# Traumatic stress history interacts with chronic peripheral inflammation to alter mitochondrial function of synaptosomes in a sex-specific manner

**DOI:** 10.1101/2020.02.12.946079

**Authors:** Gladys A. Shaw, Molly M. Hyer, Imogen Targett, Kimaya C. Council, Samya K. Dyer, Susie Turkson, Chloe M. Burns, Gretchen N. Neigh

**Author notes:** **Corresponding Author:** Gretchen N. Neigh, Department of Anatomy & Neurobiology, Virginia Commonwealth University, 1101 East Marshall Street, Box 980709, Richmond, VA 23298, Voice.

## Abstract

**Background:** Repeated exposures to chronic stress can lead to long lasting negative behavioral and metabolic outcomes. Here, we aim to determine the impact of chronic stress and chronic low-level inflammation on behavior and synaptosomal metabolism.

**Methods:** Male (n = 31) and female (n = 32) C57Bl/6 mice underwent chronic repeated predation stress or daily handling for two rounds of 15 consecutive days of exposure during the adolescent and early adult timeframes. Subsequently, mice were exposed to repeated lipopolysaccharide (LPS; 7.5 x 10^5^ EU/kg) or saline injections every third day for eight weeks. Exploratory and social behaviors were assessed in the open field and social interaction tests prior to examination of learning and memory with the Barnes Maze. Mitochondrial function and morphology were assessed in synaptosomes post-mortem. In addition, expression of TNF-α, IL-1ß, and ROMO1 were examined in the hippocampus and prefrontal cortex. Circulating pro- and anti-inflammatory cytokines in the periphery were assessed following the first and last LPS injection as well as at the time of tissue collection. Circulating ROMO1 was assessed in terminal samples.

**Results:** Exposure to repeated predatory stress increased time spent in the corners of the open field, suggestive of anxiety-like behavior, in both males and females. There were no significant group differences in the social interaction test and minimal effects were evident in the Barnes maze. A history of chronic stress interacted with chronic LPS in male mice to lead to a deficit in synaptosomal respiration. Female mice were more sensitive to both chronic stress and chronic LPS such that either a history of chronic stress or a history of chronic LPS was sufficient to disrupt synaptosomal respiration in females. Both stress and chronic LPS were sufficient to increase inflammation and reactive oxygen in males in the periphery and centrally. Females had increased markers of peripheral inflammation following acute LPS but no evidence of peripheral or central increases in inflammatory factors or reactive oxygen following chronic exposures.

**Conclusion:** Collectively, these data suggest that while metrics of inflammation and reactive oxygen are disrupted in males following chronic stress and chronic LPS, only the combined condition is sufficient to alter synaptosomal respiration. Conversely, although evidence of chronic inflammation or chronic elevation in reactive oxygen is absent, females demonstrate profound shifts in synaptosomal mitochondrial function with either a history of chronic stress or a history of chronic inflammation. These data highlight that differential mechanisms are likely in play between the sexes and suggest that female sensitivity to neurogenerative conditions may be precipitated by influence of life experiences on mitochondrial function in the synapses.

## 1. Introduction

Posttraumatic Stress Disorder (PTSD) is a unique psychiatric condition in that exposure to a life-threatening stressor is required for diagnosis (Shalev, 2009). PTSD co-occurs with inflammatory disorders and the mechanisms of this relationship are not fully defined (Neigh and Ali, 2016). In addition, metabolic disorders are more prevalent among people living with PTSD than among the general population (Lihua et al., 2020; Michopoulos et al., 2017). Furthermore, although the impact of PTSD on neural mitochondrial function is not known, PTSD is a risk factor for dementia and related disorders (Bonanni et al., 2018; Clouston et al., 2019; Desmarais et al., 2020). One critical factor in the determination of whether or not PTSD will manifest is age, with adolescents much more likely to develop PTSD compared to adults exposed to similar traumatic stressors (Van Der Kolk, 1985). In addition, repeated victimization increases risk for PTSD and comorbidities (Macdonald et al., 2010). Furthermore, repeated trauma is common; 25%-38% of adults have a history of multiple traumas (Green et al., 2000; Kessler et al., 1995), while approximately 55% of youth with a trauma history report multiple traumatic events (Macdonald et al., 2010).

Although it is not possible to model a complex neuropsychiatric condition like PTSD in a rodent, the impact of a history of life-threatening traumas can be assessed in rodent models while controlling confounding variables present in the human population (i.e. diet, tobacco, drugs, alcohol). An important area of study is the determination of the extent to which life-threatening stressors can alter the relationship between inflammatory stimuli and the neural metabolic response. Here we determined the extent to which repeated traumatic events interacted with exogenous chronic peripheral inflammation to alter neural metabolism and behavior. The use of a design that includes traumatic stressors over multiple life phases is critical because chronic developmental stress can create a susceptibility not evident until the system is again challenged in adulthood (Bourke CH et al., 2013;Pyter LM et al., 2013).

We focus on mitochondrial integrity because it is an essential component of neuronal health and function that, when compromised, leads to impaired activity, decreased synaptic connectivity, and eventual apoptosis (Adiele and Adiele, 2019; Allen et al., 2018; Smith et al., 2016; Turkson et al., 2019). Traumatic stress may translate to neural metabolic dysfunction through oxidative damage that can impair neuronal integrity (Clark et al., 2017; Fang et al., 2012) and modulate mitochondrial function (Hunter et al., 2016; Khalifa et al., 2017; Lapp et al., 2019) inducing a global change in neural metabolism (Picard et al., 2018; Turkson et al., 2019). To this end, a recent study reported metabolic signatures consistent with compromised mitochondrial function in male combat veterans, even when controlling for common confounding variables (Mellon et al., 2019). In addition, repeated shock, proposed as a model of PTSD, has been reported to cause acute alterations in mitochondrial function of hippocampal tissue of adult male rats (Seo et al., 2019).

Oxidative stress alters neuronal mitochondria trafficking (Fang et al., 2012) and can be a consequence of chronic psychological stress (for full reviews, see Bouayed et al., 2009; Salim, 2016, 2014; Smaga et al., 2015). Of particular relevance to stress-induced disorders, is evidence of sex differences in the regulation of oxidative stress, mitochondrial phenotype, and mitochondrial metabolism (Khalifa et al., 2017); however, most mitochondrial studies related to stressor exposure have focused predominantly on male subjects and measures proximate to stressor exposure. Here we report that a history of traumatic life stress interacts with both sex and chronic inflammation to alter mitochondrial bioenergetics at the presynaptic terminal. An understanding of the impact of traumatic stress on neural mitochondrial function may help to define targeted mechanistic interventions to mitigate the impact of traumatic stress on neural function.

## 2. Materials and Methods

### 2.1. Animals

Male (n = 31) and female (n = 32) C57Bl/6 mice were purchased from Taconic Laboratories and arrived at our facilities at postnatal day (PND) 22. Upon arrival, mice were pair housed with a same-sex cage-mate and housed in ventilated rack cages in a temperature and humidity-controlled room on a 12:12 light:dark cycle. Mice were given free access to food and nesting material was provided. All facilities were AAALAC approved. All animal protocols were approved by the Animal Care and Use Committee at Virginia Commonwealth University.

### 2.2. Chronic Repeated Predation Stress (CRPS)

Chronic repeated predation stress protocol was conducted as previously described and more details are available in Supplemental Methods (Burgado et al., 2014; Shaw et al., 2020). At PND 34, mice were randomly chosen to either undergo chronic repeated predation stress (CRPS; male n = 15, female n = 16) or remain in the non-stress group (male n = 16, female n = 16). CRPS occurred for 30-minutes a day over 15 consecutive days during two time periods (PND 34-48 and PND 57-71).

#### Chronic LPS Administration

Lipopolysaccharide (LPS; Sigma-Aldrich, Cat. # L4391) or saline (Hospira Inc., Lake Forest, IL, USA) was administered at a concentration of 7.5 x 10^5^ EU/kg via intraperitoneal injection (IP) to induce chronic low-level inflammation (Barnum et al., 2012). Injections were given every third day from PND 76 – 121 during the light cycle. A total of 16 injections were administered. An additional injection of LPS or saline was administered at PND 140, with subsequent physical assessments on PND 141 to assess the somatic reaction to an immunological challenge and any acute changes in cognition (via Barnes Maze). Assessments of weight (g), mobility (graded scale: 1 = normal mobility, 2 = mobile if disturbed, and 3 = immobile), and piloerection (graded scale, zero = none, 0.5 = only around neckline, 1 = full coat) were taken 24 and 48 hours after each injection.

### 2.3. Plasma Collection

In addition to terminal samples, minimum volume blood samples were collected via retro-orbital bleeds at three time points: two hours after the first injection (PND 76), two hours after the 16^th^ injection (PND 121), and two hours after the LPS trigger injection (PND 141). For retro-orbital samples, mice were anesthetized via isoflurane inhalation and blood collected using a 100µL capillary tube. Approximately 50µL of blood was collected from each retro-orbital bleed. Samples were placed in two-mL EDTA coated tubes and spun at 1800xg for 20 minutes. Terminal blood samples were collected following rapid decapitation into EDTA coated tubes. Plasma was isolated, placed in an eight-tube strip and stored at −80°C until further use.

### 2.4. Behavioral Assessments

To assess exploratory and anxiety-like behavior (Burgado et al., 2014; Carola et al., 2002; Choleris et al., 2001; Prut and Belzung, 2003), all mice were subject to the open field test the morning before their eighth injection (PND 97), 72 hours after the seventh LPS exposure. Mice were permitted ten minutes of exploration in the open field during their light cycle. Behaviors were recorded using an overhead camera and EthoVision XT 14.0 software (Noldus Technologies, Leesburg, VA, USA). Time spent in the center of the arena, as well as velocity and distance traveled, were assessed via overhead camera and EthoVision XF 14.0. All female mice underwent a visual assessment for the estrus cycle immediately following assessments. Visual characterization used were based on those found in Byers et al., 2012. To maintain consistency between sexes, males were handled in a similar manner at the same time of day. No more than two females in any one group were in estrous at any data collection point so this was not included as a factor in subsequent analyses. Social interaction and learning and memory via the Barnes Maze were also assessed. Methods for these assays are presented in the Supplemental Methods.

### 2.5. Tissue Collection

Mice were euthanized via rapid decapitation on PND 147-148. This timepoint is seven to eight days following the final LPS trigger injection. Samples had to be collected across two days due to the temporal dynamics of the mitochondrial assay. All groups were balanced between collection days. Additional details on synaptosomal isolation are available in Supplemental Methods.

### 2.6. Mitochondrial Respiration

Mitochondrial respiration was measured using Agilent’s Cell Mitochondrial Stress Test kit (Agilent Technologies, PN: 103015-100) and Seahorse XFe24 Analyzer. Drug injections were prepared according to manufacturer instructions at 2.0µM Oligomycin, 1.0µM FCCP, and 0.5µM Rotenone/Antimycin A. Oxygen consumption rates (OCR) were determined by sequential measurement cycles consisting of a 30 second mixing time followed by a two-minute wait time and then a three-minute measurement period (three measurements following each reagent addition). Reagents were added to SeahorseXFe24 FluxPak in dilutions according to manufacturer’s recommendation (2.0μM Oligomycin, 1.0μM carbonyl cyanide-p-trifluoromethox- yphenyl-hydrazon (FCCP), 0.5μM Rotenone/antimycin A per well). The first three measurements of OCR occur prior to addition of mitochondrial reagents and indicates basal respiration. Oligomycin is a complex V inhibitor and OCR following this addition indicates ATP-linked respiration (subtraction of baseline OCR) and proton leak respiration (subtraction of non-mitochondrial respiration). FCCP is a protonophore and adding it will collapse the inner membrane gradient and push the electron transport chair to the maximal rate. Inhibition of complex III and I is achieved through addition of antimycin A and rotenone which will terminate electron transport chain function and demonstrate non-mitochondrial respiration. Mitochondrial reserve is calculated by subtraction of basal respiration from maximal respiration (Rose et al., 2014). Data were recorded in Agilent’s Wave Software and subsequently exported via Excel document for further analysis.

### 2.7. Western Blots

Western blots were used to assess the relative mitochondrial content from the synaptosomal isolations. Antibodies for presynaptic terminals (SNAP-25; ThermoFisher Scientific, Waltham, MA; Cat. # PA1-9102), and mitochondria (Hexokinase I; ThermoFisher Scientific, Cat. # MA5-15680) assess mitochondrial concentration and relative presence of presynaptic terminals within synaptosomal preparations. Additional details are available in Supplemental Methods.

### 2.8 Mitochondrial Assessment

Following Transmission Electron Microscopy (TEM; see Supplemental Methods for procedural details), mitochondrial phenotype was assessed using the Flaming scoring system (Flameng et al., 1980; Hawong et al., 2015; Li et al., 2018). Mitochondrial phenotype was scored on a scale of one to five, with five signifying healthy mitochondria, four signifying agranular mitochondria, three signifying inflamed mitochondria with intact membranes and cristae, two signifying mitochondria with intact membranes but broken cristae, and one signifying mitochondria with broken membranes and cristae. Scores were assessed using TEM images taken at 4000x magnification by a blind counter.

### 2.8. Peripheral Inflammation

Plasma samples collected from three time points: 1) two hours after the first chronic injection (acute), 2) two hours after the final chronic injection (chronic), and 3) from trunk blood at the time of tissue collection (terminal; seven to eight days after a trigger LPS injection used for cognitive testing) to assess markers of peripheral inflammation. Circulating concentrations of IFN-γ, IL-1ß, IL-2, IL-4, IL-5, IL-6, IL-10, IL-12p70, KC/GRO, and TNF-α were assessed using the MSD V-Plex Proinflammatory Panel 1 Mouse Kit (Meso Scale Diagnostics, Rockville, MD, USA; Cat. #: K15048D-2) and read using the MESO QuickPlex SQ 120 (Meso Scale Diagnostics). A standard curve was completed to determine the required dilutions for analysis. Based on results from the standard curve, samples from saline treated mice were diluted 1:8 for both the acute and chronic time points. Samples from LPS treated mice were diluted 1:20 and 1:30 for acute and chronic time-points respectively. All terminal time points regardless of treatment history were diluted 1:2. MSD workflow was completed according to manufacturer instructions.

### 2.9. ROMO-1 ELISA

Circulating ROS was assessed via enzyme linked immunosorbent assay (ELISA) for ROMO1 (MyBioSource, San Diego, CA; Cat # MBS3805476) on plasma isolated from trunk blood samples. All plasma samples were diluted 1:5 for analysis in sample diluent as suggested by the manufacturer. ELISA workflow was completed according to manufacturer’s instructions.

### 2.10. Quantitative Polymerase Chain Reaction (qPCR)

Relative transcript levels of the inflammatory cytokines IL-1ß (ThermoFisher Scientific, Cat. # Mm00434228_m1), and TNFα (Cat. # Mm00443258_m1), and the reactive oxygen species modulator ROMO1 (Cat. # Mm01246687_g1), were assessed via TaqMan qPCR in both the hippocampus and prefrontal cortex of each mouse. Two housekeeping genes, GAPDH (Cat. # Mm99999915_g1), and ß-actin (Cat. # Mm01205647_g1) were used for normalization of each sample. All samples were run in triplicate.

### 2.11. Statistical Analysis

All statistical data was analyzed within each sex using GraphPad Prism 8.2.0 for Windows (GraphPad Software, La Jolla, CA). Physical Weights, behavioral data from the open field, and qPCR were analyzed using two-way analysis of variance (ANOVA) with the factors of stress and LPS. Piloerrection and immobility were measured using a generalized linear mixed-model. Synaptosomal respiration data was normalized according to previously published methods (Choi et al., 2009) such that non-mitochondrial respiration (measurements ten through twelve) were set to an average OCR of zero for each subject. Normalized OCR data was analyzed using a three-way ANOVA with the factors stress, LPS, and measurement. Western blot data was analyzed using a two-way ANOVA with the factors stress and LPS. The Flameng score and MSD data were analyzed by two-way ANOVAs with the factors of stress and LPS. TaqMan PCR data were analyzed using a two-way ANOVA of the 2^ΔΔ^ct values with the factors stress and LPS. ROMO1 ELISA data was analyzed using a two-way ANOVA with the factors of stress and LPS. Peripheral cytokine data were analyzed using a three-way ANOVA with the factors stress, LPS, and time point. For all three-way ANOVAs, when significant interactions were observed, two-way ANOVAs and appropriate post hoc comparisons were conducted to determine factors driving group differences. All statistical tests were run with a significance level (α) set to 0.05.

## 3. Results

### 3.1. Physical Assessments

Body mass increased over the duration of the study for male subjects regardless of stress or LPS exposure (F_(1,27)_ = 63.87, p < 0.0001; **FIGURE 1A**). However, there were timepoint specific group differences in body mass (F_(1,27)_ = 10.84, p = 0.0028). Specifically, body mass of males treated with LPS was lower than saline injected counterparts following the first exposure to LPS (p = 0.0225). Mobility was similarly impacted such that mobility increased over time (F_(1,27)_ = 5.841, p = 0.0227), but was differentially impacted by LPS versus saline treatment (F_(1,27)_ = 5.841, p = 0.0227) in a time-specific manner (F_(1,27)_ = 5.841, p = 0.0227). Post hoc analysis isolated this effect to the initial LPS exposure (p = 0.0019; **FIGURE 1C**), with males treated with saline showing a higher mobility score than males treated with LPS. Last, in terms of piloerection, males exhibit a main effect of time (F_(1,27)_ = 9.833, p = 0.0041), stress (F_(1,27)_ = 5.849, p = 0.0226), and treatment (F_(1,27)_ = 6.692, p = 0.0154; **FIGURE 1E**) with no interactions such that both LPS and chronic stress precipitated increased piloerection over time.

**Figure 1:**
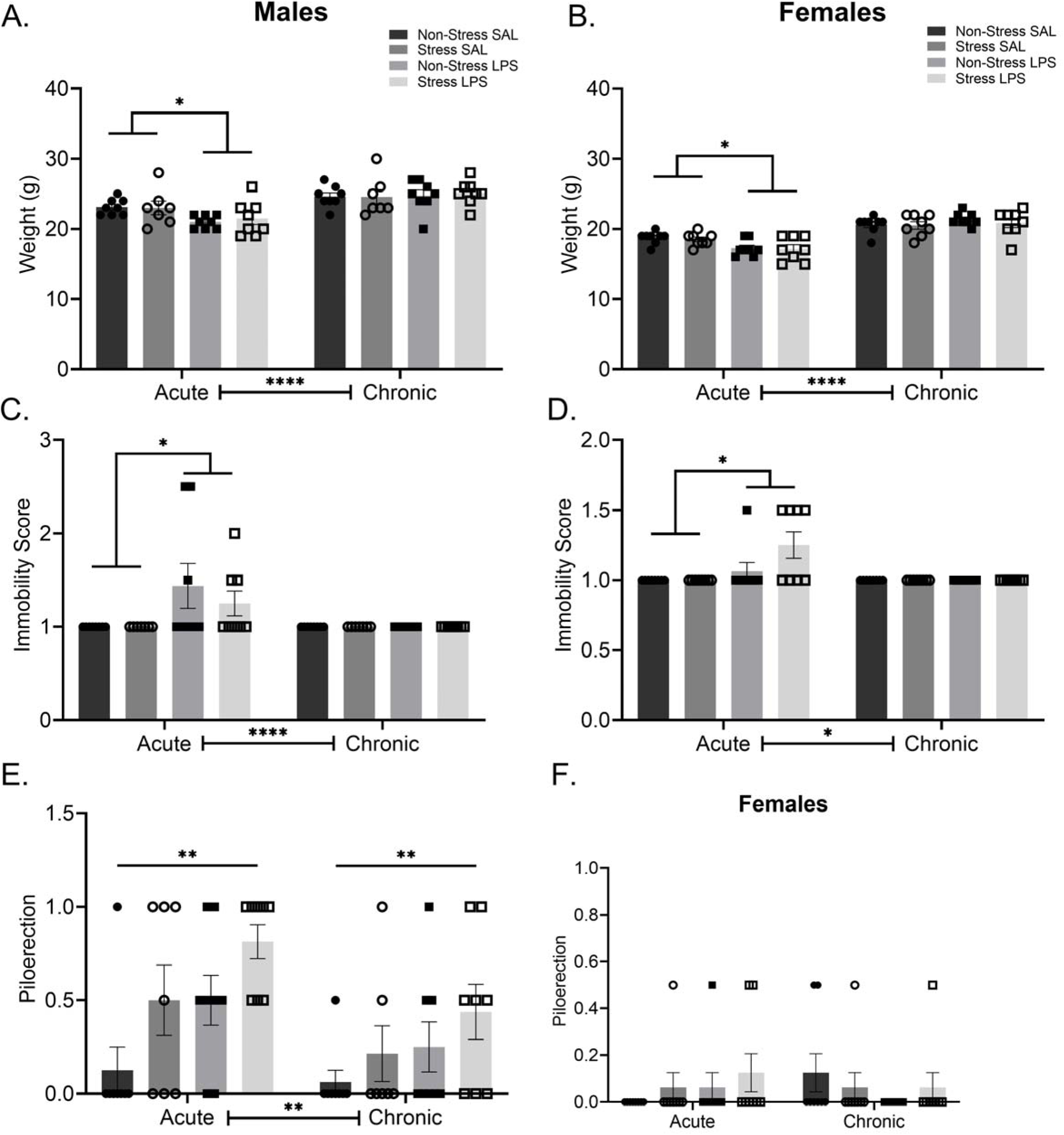
Physical Changes Following Chronic Stress and Chronic LPS. **A)** Acute, but not chronic, LPS reduced body mass in male and, **B)** female mice. **C)** Acute LPS reduced mobility in males but this reduction was no longer evident after chronic LPS exposure. **D)** LPS treated females also show decreased mobility 24 hours following the initial treatment injection which was no longer present at the final injection. **E)** Piloerection was elevated in males following either LPS or chronic stress. **F)** Female piloerection showed no significant differences at either the acute of chronic treatment time points. Reported values depict mean ± SEM. *p < 0.05, **p < 0.01, ****p < 0.0001.

Within the females, body mass increased over time (F_(1,28)_ = 219.3, p < 0.0001) and this effect was modified by exposure to LPS interaction (F_(1,28)_ = 25.42, p < 0.0001; **FIGURE 1B**). Post hoc analysis identified that, like the males, LPS treated females had lower body mass following the first LPS injection than did saline exposed controls (p = 0.0051). Mobility scores differed as a function of time (F_(1,28)_ = 7.609, p = 0.0101), LPS treatment (F_(1,28)_ = 7.609, p = 0.0101) with evidence of an interaction (F_(1,28)_ = 7.609, p = 0.0101; **FIGURE 1D**). This difference was specific to decreased mobility after the first LPS injection compared to saline injected controls (p = 0.0010). Piloerection differed by time and LPS exposure (F_(1,28)_ = 4.308, p = 0.0472; **FIGURE 1F**), but post hoc analysis did not isolate this difference to a specific set of comparisons (p > 0.05).

### 3.2. Open Field

Time in the center of the open field was not different among males regardless of a history of stress or LPS exposure (p > 0.05; **FIGURE 2A**). A history of stress exposure did lead to increased time in the corners of the open field regardless of LPS exposure (F_(1,27)_ = 5.857, p = 0.0225; **FIGURE 2B**). Female mice with a history of stress exposure spend less time in the center of the open field than controls, regardless of LPS exposure (F_(1,28)_ = 4.166, p = 0.0508). In addition, a history of stress caused an increase in the time spent in the corners of the arena for females (F_(1,28)_ = 7.256, p = 0.0118). Distance traveled in the open field was increased for both male and female mice with a history of chronic stress as compared to their respective controls (males: F_(1,27)_ = 7.413, p = 0.0112; females: F_(1,28)_ = 7.628, p = 0.0100; **FIGURE 2C**).

**Figure 2:**
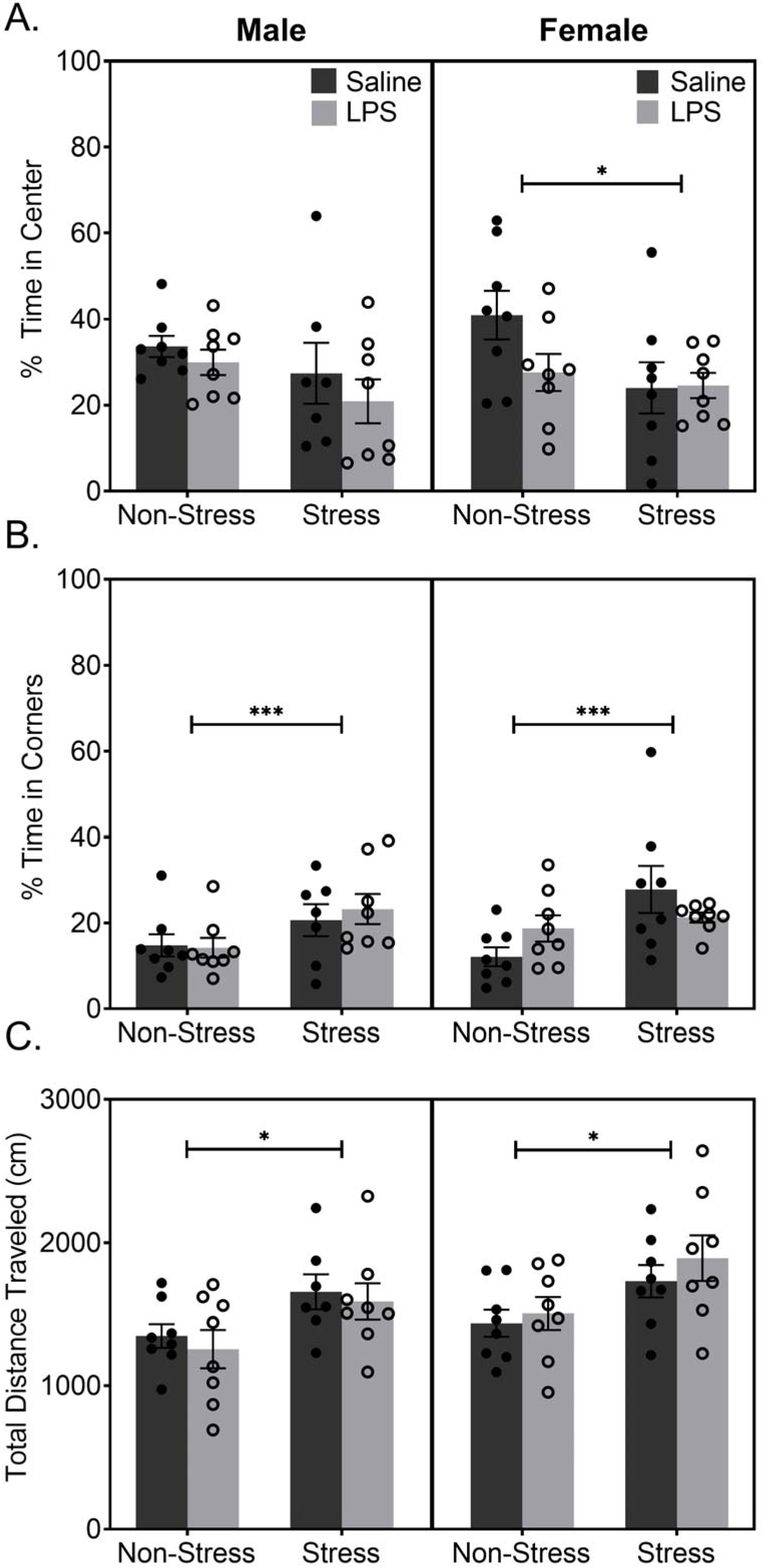
A History of Stress Increases Anxiety-Like Behaviors in the Open Field. **A)** Male data show no significant difference in the percent of time spent in the center of the arena. Female data show a main effect of stress in time spent in the center of the arena, with females with a history of stress spending less time in the center suggesting increased anxiety-like behavior. **B)** Both males and females show a significant difference in time spent in the corners of the arena, with mice with a history of stress spending increased time in the corners. **C)** Analysis of movement patterns suggested mice with a history of stress, regardless of sex, appear to be more hyperactive than their sex matched controls. Reported values depict mean ± SEM. *p < 0.05, ***p < 0.001.

### 3.3. Synaptosomal Respiration

Although a history of chronic stress did not alter mitochondrial synaptosomal respiration in samples from male rats (p > 0.05), synaptosomal respiration was altered by chronic LPS treatment within males (F_(1,276)_ = 13.72, p = 0.0003). In addition, a history of chronic stress interacted with chronic LPS to further exacerbate the impact of LPS alone (F_(1,276)_ = 3.959, p = 0.0476; **FIGURE 3A**). Specifically, significant group differences existed between LPS treated males with no stress history and LPS treated males with a history of stress (p = 0.0011) and between saline treated males with no stress history and LPS treated males with no stress history (p < 0.0001). This indicates that the LPS treated males with a history of stress had a significant decrease in overall OCR when compared to all other groups. A breakdown of each individual component of the OCR waveform showed no significant differences in basal respiration (p > 0.05, **FIGURE 3B**), maximal respiration (p > 0.05, **FIGURE 3C**), proton leak (p > 0.05, **FIGURE 3D**), ATP production (p > 0.05, **FIGURE 3E**), or spare respiratory capacity (p > 0.05, **FIGURE 3F**). Collectively, these results indicated an overall impact on mitochondrial respiration that is not specific to any one part of the electron transport chain.

**Figure 3:**
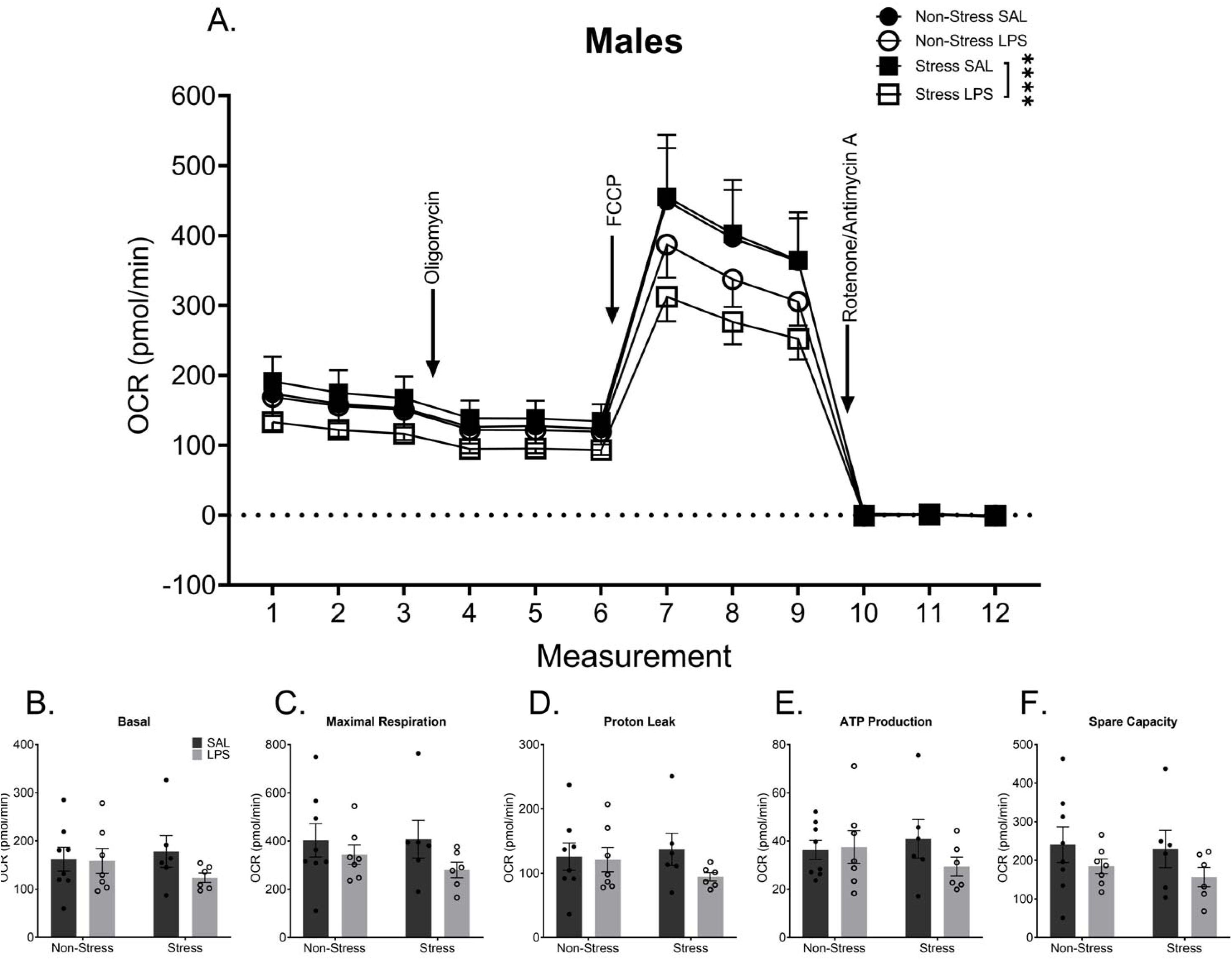
Chronic Stress Combined with Chronic LPS Alters Synaptosomal Respiration in Male Mice. **A)** Metabolic data display a significant decrease in synaptosomal respiration in males with a history of stress and LPS treatment when compared to males with a history of stress alone. Despite this overall difference, there are no significant differences in basal respiration **(B),** maximal respiration **(C),** proton leak (**D),** ATP production (**E),** or spare capacity **(F).** Reported values depict mean ± SEM. ****p < 0.0001.

Mitochondrial synaptosome respiration was altered by a history of chronic life stress (**FIGURE 4A**) or a history of chronic LPS treatment (F_(1,300)_ = 5.725, p = 0.0173) or the combination of the two (F_(1,300)_ = 33.16, p < 0.0001). Furthermore, differences were somewhat dependent on which aspect of the electron transport chain was being probed, as evidenced by a measurement by LPS treatment by stress interaction (F_(11,300)_ = 2.269, p = 0.0114). Measurement refers to timepoints in the mitochondrial assay and which portion of the electron transport chain is being inhibited or stimulated. A history of chronic life stress increased mitochondrial respiration as compared to controls without stress exposure (p = 0.0002). Chronic LPS treatment also increased mitochondrial respiration as compared to saline treated controls, when no history of stress was present (p = 0.0104). When both a history of chronic stress and chronic LPS were present, synaptosomal respiration was reduced compared to stress alone (p < 0.0001) or LPS alone (p < 0.0001), but this group was not different than controls (p > 0.05). Isolation of each individual component of the OCR waveform show a significant difference in proton leak within females with a history of stress (p = 0.0468), with saline treated females exhibiting decreased proton leakage compared to females that were exposed to both chronic stress and chronic LPS (**FIGURE 4D**). There were no significant differences in basal respiration (p > 0.05, **FIGURE 4B**), maximal respiration (p > 0.05, **FIGURE 4C**), ATP production (p > 0.05, **FIGURE 4E**), or spare respiratory capacity (p > 0.05, **FIGURE 4F**).

**Figure 4:**
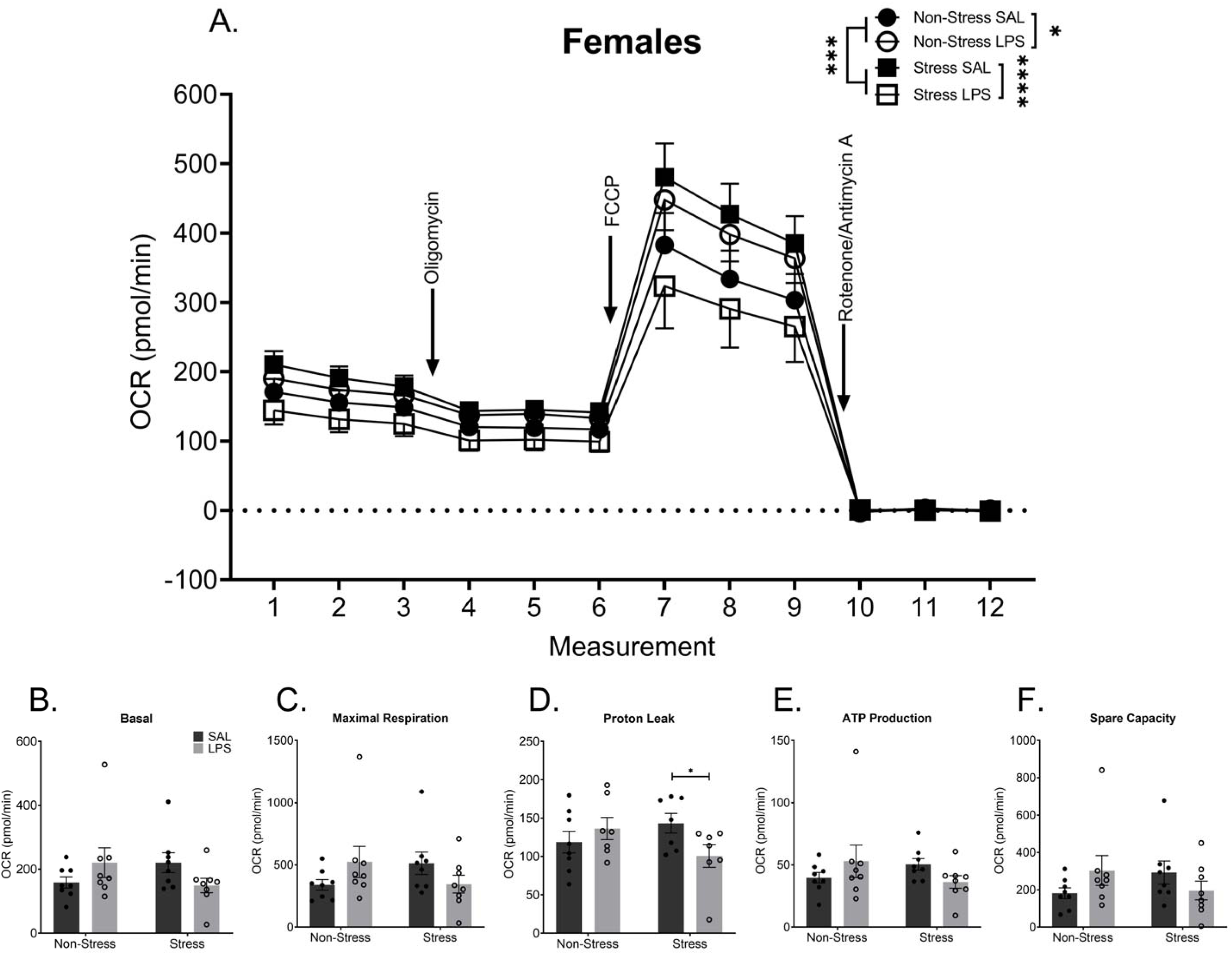
Chronic Stress and Chronic LPS Independently Alter Synaptosomal Respiration in Female Mice. **A)** Metabolic data display a stress and treatment dependent change in overall synaptosomal respiration. Chronic stress increased synaptosomal respiration even 11 weeks removed from the final stressor. In addition, chronic LPS treatment increased overall synaptosomal respiration in mice with no history of stress. Conversely, females that had both a history of chronic stress and were exposed to chronic LPS treatment demonstrated reduced overall synaptosomal respiration compared to the other groups. Although effects were not specific to basal respiration **(B),** maximal respiration **(C),** ATP production (**E),** or spare capacity **(F)**, the combination of stress and LPS caused a decrease in proton leak **(D)** in females. Reported values depict mean ± SEM. *p < 0.05, ****p < 0.0001.

### 3.4. Western Blot

Pooled synaptosomal preparations were probed for mitochondria (via Hexokinase I) and pre-synaptic terminals (via SNAP-25; **SUPPLEMENTARY FIGURE 3**). Within both males and females, there was no significant fold change difference in mitochondrial presence as a function of either stress or LPS treatment, as measured via Hexokinase I compared to same sex controls (p > 0.05; **FIGURE 5A-B**). Within the males, there was no significant difference in pre-synaptic terminal presence due to either stress or LPS (p > 0.05; **FIGURE 5A**). Conversely, within the females, data show a significant interaction among stress history and LPS exposure (F_(6,24)_ = 3.421, p = 0.0139). Post hoc analysis demonstrated that LPS treatment, regardless of stress history, led to an increase in pre-synaptic terminal presence as compared to controls (p = 0.0214 and p = 0.0168 respectively; **FIGURE 5B**).

**Figure 5:**
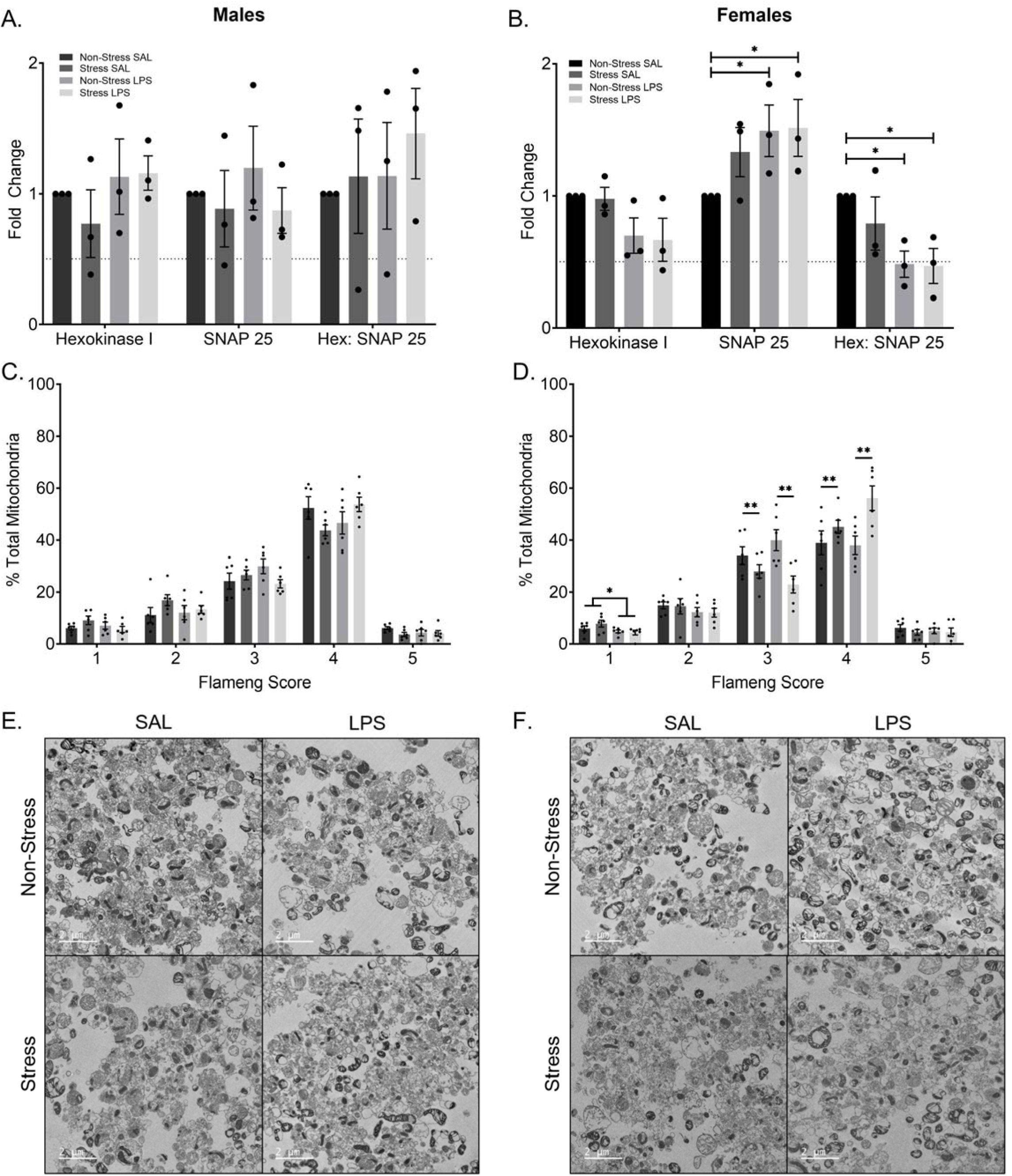
Female Mice Display Synaptosomal Composition and Mitochondrial Phenotype Differences. **A)** There is no significant difference in the number of mitochondria or presynaptic terminals in male mice. Likewise, there is no difference in relative number of mitochondria per presynaptic terminal. **B)** Within the females, there is no significant difference in mitochondrial number. There is a significant increase in the presence of presynaptic terminals in the LPS treated groups when compared to non-stress saline treated controls. Moreover, there is a significant decrease in the relative number of mitochondria per presynaptic terminal in the LPS treated groups when compared to the non-stress saline treated controls. **C)** Flameng score, a measure of mitochondrial phenotype and proxy for mitochondrial function did not differ by group for males. **D)** Flameng scores within the females show those who endured chronic LPS treatment display decreased mitochondria with broken membranes and broken cristae (Flameng score of one). A history of stress decreases inflamed mitochondria (Flameng score of three) and increases agranular mitochondria (Flameng score of four) within the female mice regardless of LPS exposure. Representative transmission electron images of males **(E)** and females **(F).** Reported values depict mean ± SEM. *p < 0.05, **p < 0.01.

The ratio of Hexokinase I to SNAP-25 was calculated to assess the relative amount of mitochondria per presynaptic terminal. Males did not differ in relative ratio as a function of either stress or LPS exposure (p > 0.05). Females treated with either LPS alone (p = 0.0159) or with the combined exposure to stress and LPS (p = 0.0136) had a lower ratio indicative of fewer mitochondria per pre-synaptic terminal as compared to controls (no stress, no LPS).

### 3.5. Phenotypic Mitochondrial Assessment

In the males, there were no significant differences based on sex or treatment in the relative percentage of mitochondria at any Flameng score (p > 0.05; **FIGURE 5C**). Representative TEM images for each male group are displayed in **FIGURE 5E**. Female data show a main effect of LPS treatment in mitochondria with broken membranes and cristae (Flameng score of one; F_(1,20)_ = 5.601, p = 0.0281; **FIGURE 5D**) and a main effect of stress in both inflamed mitochondria (Flameng score of three; F_(1,20)_ = 11.73, p = 0.0027) and agranular mitochondria (Flameng score of four; F_(1,20)_ = 9.466, p = 0.0060). Representative TEM images for each female group are displayed in **FIGURE 5F**.

### 3.6 qPCR

Chronic LPS exposure increased expression of IL-1ß in the hippocampus (F_(1,27)_ = 11.12, p = 0.0025; **FIGURE 6A**) of male mice. In addition, both chronic LPS (F_(1,27)_ = 5.176, p = 0.0311) and chronic stress (F_(1,27)_ = 5.539, p = 0.0261) increased TNFα expression in the male hippocampus. No effects of chronic stress or chronic LPS were detected in the prefrontal cortex in expression of either TNFα (p > 0.05) or IL-1ß (p > 0.05). No impact of either a history of chronic stress or a history of chronic LPS was observed in terms of expression of IL-1ß or TNFα in the hippocampus or prefrontal cortex of female mice (p > 0.05; **FIGURE 6B**). Likewise, ROMO1 expression did not differ by either stress or LPS history for either males or females (p > 0.05).

**Figure 6:**
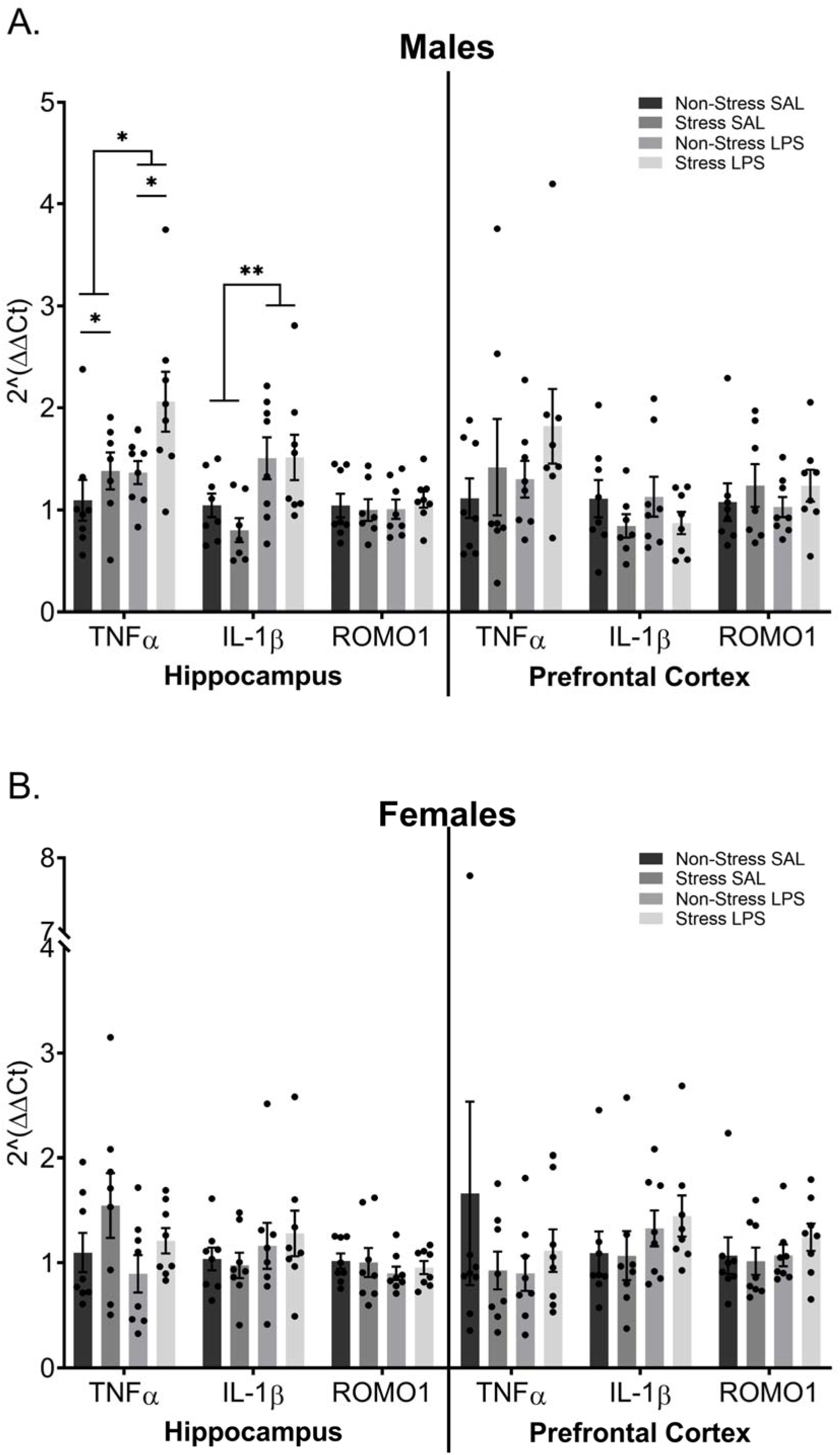
Chronic Inflammation and a History of Chronic Stress Alter Expression of TNF-α and IL-1ß in the Hippocampus of Male Mice. Transcript levels of the pro-inflammatory cytokines *TNF-α* and *IL-1ß* as well as the reactive oxygen species marker *ROMO1* were assessed in hippocampal and prefrontal cortex tissue via TaqMan RT-qPCR. **A)** *TNF-α* levels within male hippocampi show that both stress and LPS increase *TNF-α,* yet display no significant differences in the prefrontal cortex. Chronic LPS treatment increased *IL-1ß* transcript in the hippocampus of male mice, but did not change transcript levels in the prefrontal cortex. Neither hippocampal not prefrontal cortex samples showed significant differences in levels of *ROMO1*. **B)** Female data show no significant differences in *TNF-α, IL-1ß,* or *ROMO1* levels in either the hippocampus or prefrontal cortex. Reported values depict mean ± SEM. *p < 0.05, **p < 0.01.

### 3.6. Peripheral Inflammation

#### 3.6.1. Proinflammatory Cytokines

TNFα concentrations in male serum differed by sample collection timepoint (F_(2,77)_ = 22.92, p < 0.0001), LPS treatment (F_(1,77)_ = 25.66, p < 0.0001), and stress exposure history (F_(1,77)_ = 4.392, p = 0.0394). In addition, interactions occurred between study time point and LPS treatment (F_(2,77)_ = 22.77, p < 0.0001), study time point and stress history (F_(2,77)_ = 4.603, p = 0.0129), LPS treatment and stress history (F_(1,77)_ = 4.364, p = 0.0400), and LPS treatment by study time point by stress history (F_(2,77)_ = 4.581, p = 0.0132; **FIGURE 7A**). Post hoc analysis demonstrated that males given an acute LPS injection had elevated TNFα concentrations regardless of stress history (p’s < 0.05). TNFα concentrations in females were altered by study time point (F_(2,78)_ = 31.49, p < 0.0001) and LPS treatment (F_(1,78)_ = 35.20, p < 0.0001), with a time point by LPSP treatment interaction (F_(1,77)_ = 31.43, p < 0.0001; **FIGURE 7B**). Post hoc analysis exposes this interaction is driven by a significant difference between saline and LPS treated females at the acute time point (p < 0.0001).

**Figure 7:**
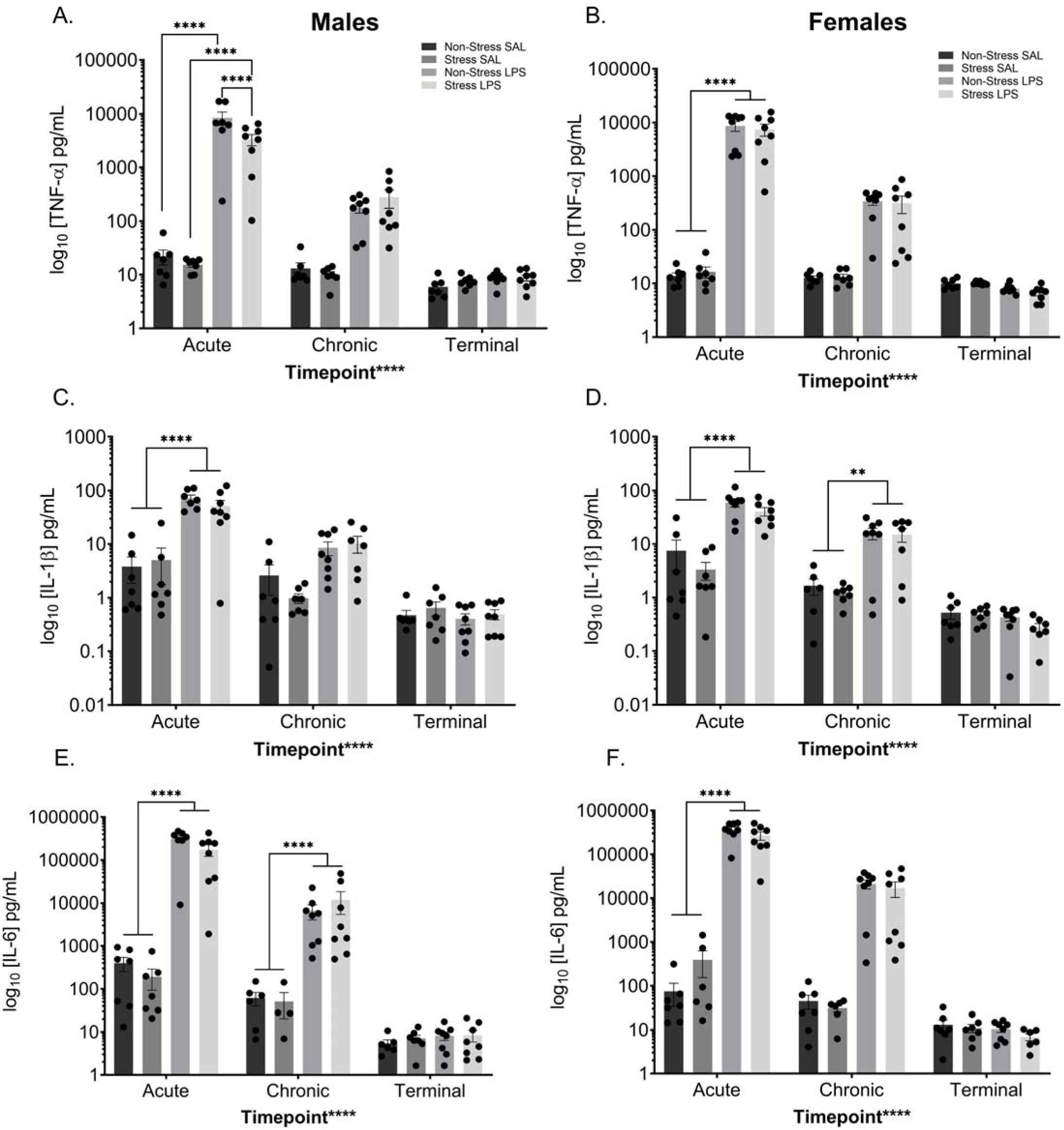
Peripheral Pro-inflammatory Cytokines Changes Following LPS. Peripheral cytokine levels were assessed at three time points; two hours after the initial treatment injection (“acute”), two hours after the last treatment injection (“chronic”), and from terminal collections (“terminal”). Chronic LPS injections increased circulating TNF-α in both males **(A**) and females (**B)** at the acute time point only. Moreover, in males, the combination of stress and LPS decreased circulating TNF-α levels when compared to those treated with LPS alone at the acute time point. Peripheral IL-1ß levels increased with chronic LPS injections at the acute time point of both males (**C)** and females (**D).** Additionally, chronic LPS treatment increased circulating IL-1ß levels in females at the chronic time point (**D**). IL-6 levels in the periphery were significantly increased following LPS treatment at both the acute and chronic time point in males (**E**). In females, IL-6 levels significantly increased following LPS treatment at the acute time point only (**F**). Reported values depict mean ± SEM. **p < 0.01, ****p < 0.0001.

Concentrations of circulating IL-1ß in males differed as a function of study time point (F_(2,76)_ = 40.39, p < 0.0001) and LPS treatment (F_(1,76)_ = 46.09, p < 0.0001; **FIGURE 7C**). Further, there is a time point by LPS treatment interaction (F_(2,76)_ = 31.82, p < 0.0001), which is driven by a difference between saline and LPS treated males at the acute time point (p < 0.0001). IL-1ß levels in the females demonstrates a main effect of study time point (F_(2,75)_ = 336.93, p < 0.0001), LPS treatment (F_(1,75)_ = 52.04, p < 0.0001), and a time point by treatment interaction (F_(2,75)_ = 24.58, p < 0.0001; **FIGURE 7D**) driven by a significant difference between saline and LPS treated females at both the acute (p < 0.0001) and chronic (p = 0.0053) study time points.

Both male and female data for IL-6 have similar patterns compared to their respective controls, with both sexes displaying main effects of study time point (male: F_(2,72)_ = 33.68, p < 0.0001; **FIGURE 7E**; female F_(2,74)_ = 50.69, p < 0.0001; **FIGURE 7F**) and LPS treatment (male: F_(1,72)_ = 36.00, p < 0.0001; female: F_(1,74)_ = 59.82, p < 0.0001), with a time point by LPS treatment interaction (males: F_(2,72)_ = 33.53, p < 0.0001; females: F_(2,74)_ = 50.55, p < 0.0001). In both cases, this interaction is driven by differences between the saline and LPS treated mice at the acute LPS time point (males: p < 0.0001; females: p < 0.0001)

Although male IFN-γ gave rise to no significant results (**FIGURE 8A**), female data showed a main effect of time point (F_(2,70)_ = 3.614, p = 0.0321) and a time point by LPS treatment interaction (F_(2,70)_ = 15.52, p < 0.0001; **FIGURE 8B**). Post hoc analysis revealed this interaction is driven by a difference between the saline and LPS treated females at the acute (p < 0.0001) and terminal (p = 0.0046) time points. This is notable as it suggests that chronic low-level inflammation increases IFN-γ circulation following an acute injection but gives rise to decreased IFN-γ circulation following chronic LPS injections.

**Figure 8:**
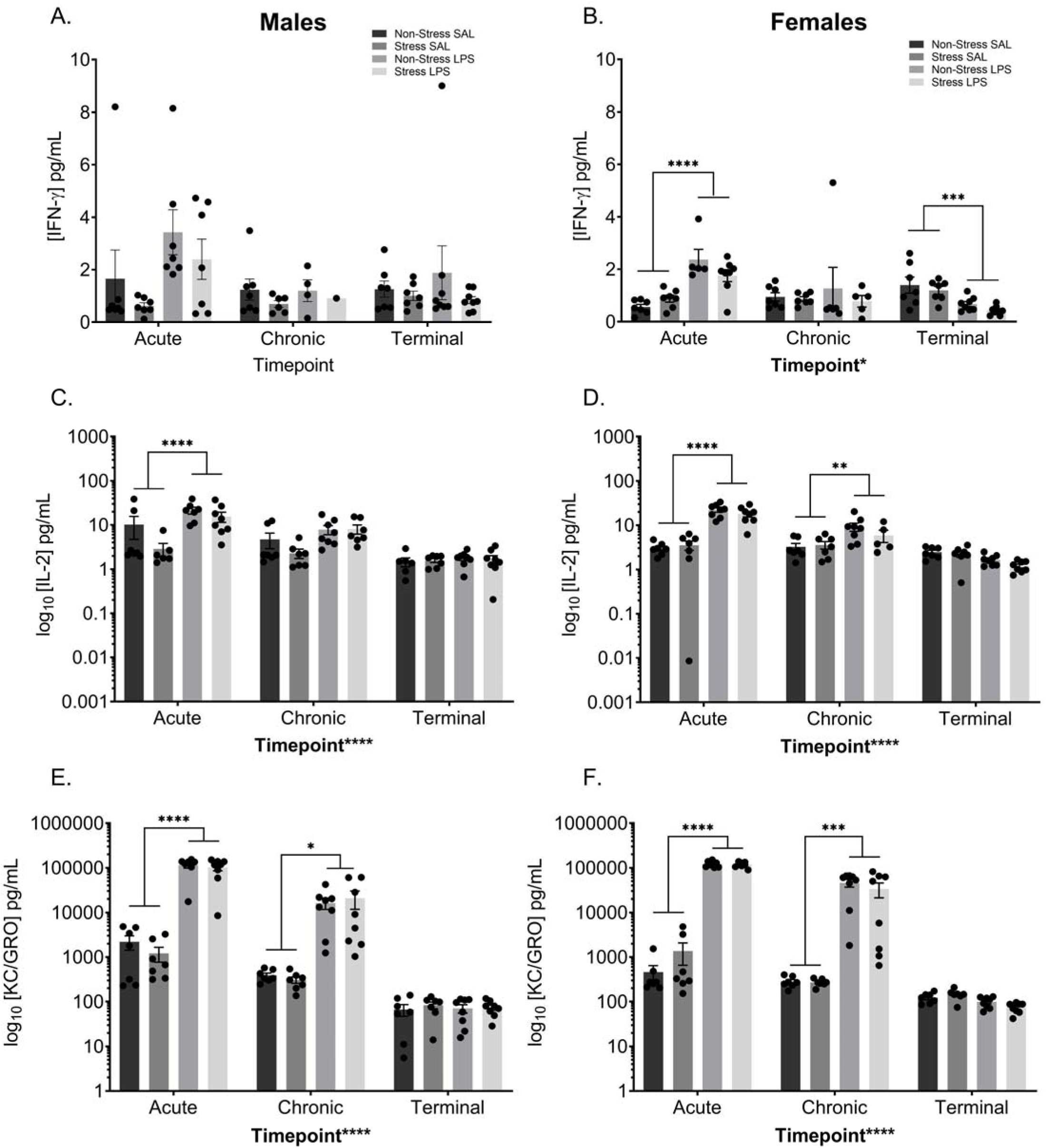
Peripheral Pro-inflammatory Cytokines Altered Following Chronic Stress and LPS. Peripheral cytokine levels were assessed at three time points; two hours after the initial treatment injection (“acute”), two hours after the last treatment injection (“chronic”), and from terminal collections (“terminal”). **A)** There was no significant difference in IFN-γ in the males. **B)** In the females, LPS treatment significantly increased circulating IFN-γ levels at the acute time point. Alternatively, a history of chronic LPS decreased baseline IFN-γ levels in the periphery of female mice. **C)** LPS significantly increased circulating IL-2 levels in male mice at the acute time point. **D)** In females, LPS increased circulating IL-2 levels at the acute and chronic time points. At both the acute and chronic times, LPS increased circulating levels of KC/GRO in both male **(E)** and female **(F)** mice. Reported values depict mean ± SEM. *p < 0.05, **p < 0.01, ***p < 0.001, ****p < 0.0001.

Proinflammatory cytokine IL-2 also show similar results within the sexes, with males and females displaying main effects of time point (male: F_(2,74)_ = 19.14, p < 0.0001; **FIGURE 8C**; female F_(2,75)_ = 49.71, p < 0.0001; **FIGURE 8D**) and LPS treatment (male: F_(1,74)_ = 14.80, p = 0.0003; female: F_(1,75)_ = 65.01, p < 0.0001), with a time point by treatment interaction (males: F_(2,74)_ = 5.650, p = 0.0052; females: F_(2,75)_ = 42.24, p < 0.0001). Within the males, the interaction is driven by a significant difference in treatment at the acute time point (p < 0.0001). Female data display a more robust change, with the interaction driven by treatment differences at both the acute (p < 0.0001) and chronic (p = 0.0033) time points.

KC/GRO, a proinflammatory cytokine that has recently been tagged as a potential biomarker of depression in the elderly (Fanelli et al., 2019) shows similar results between males and females. Data show main effects of time point (male: F_(2,77)_ = 56.34, p < 0.0001; **FIGURE 8E**; female F_(2,78)_ = 128.9, p < 0.0001; **FIGURE 8F**) and treatment (male: F_(1,77)_ = 85.65, p < 0.0001; female: F_(1,78)_ = 284.7, p < 0.0001), with a time point by treatment interaction (males: F_(2,77)_ = 53.05, p < 0.0001; females: F_(2,78)_ = 125.5, p < 0.0001). In both this interaction is driven by significant differences between saline and LPS treated mice at the acute (male: p < 0.0001; female: p < 0.0001) and chronic (male: p = 0.0177; female: p < 0.0001) time points.

IL-12p70 was below the limit of detection for most samples. A summary of the data collected can be found in **SUPPLEMENTARY TABLE 1**.

#### 3.6.2. Anti-inflammatory Cytokines

The anti-inflammatory cytokines IL-4, IL-5, and IL-10 were assessed and analyzed identically to the proinflammatory cytokines. IL-4 levels in the males display a main effect of time point (F_(2,40)_ = 13.69, p < 0.0001), treatment (F_(1,40)_ = 7.775, p = 0.0081), and a time point by treatment interaction (F_(2,40)_ = 4.207, p = 0.0220; **FIGURE 9A**). At the acute time point, there is a significant difference between the saline treated and LPS treated males (p = 0.0001), driving this interaction. Female IL-4 levels display a main effect of time point (F_(2,45)_ = 10.40, p = 0.0002) and a time point by treatment interaction (F_(2,45)_ = 6.957, p = 0.0023; **FIGURE 9B**). Similar to the males, the female interaction is driven by a significant difference between the saline and LPS treated groups at the acute time point (p < 0.0001).

**Figure 9:**
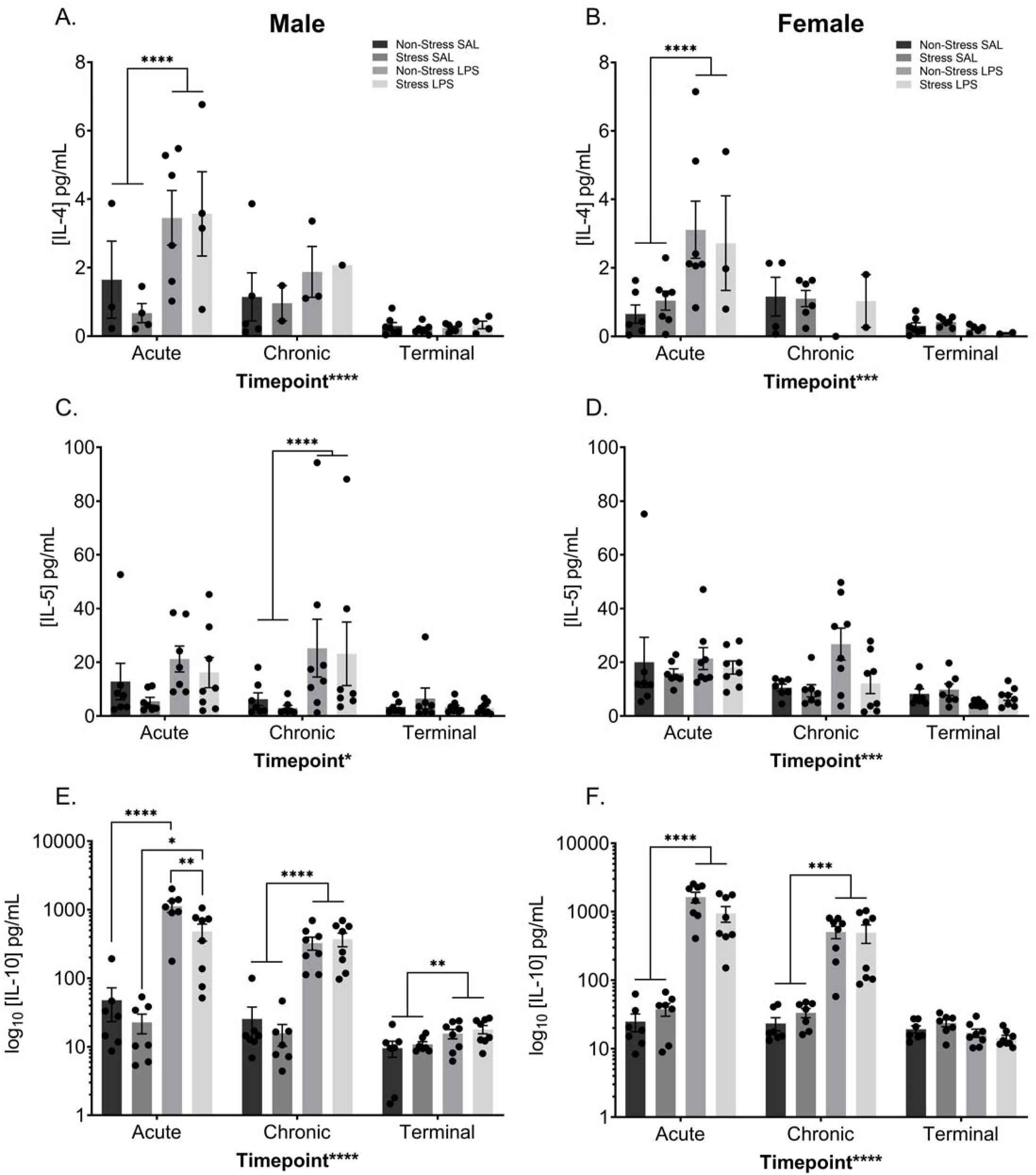
Peripheral Anti-inflammatory Cytokines Increase Following LPS. Peripheral cytokine levels were assessed at three time points; two hours after the initial treatment injection (“acute”), two hours after the last treatment injection (“chronic”), and from terminal collections (“terminal”). LPS treatment increased circulating IL-4 levels in both males **(A)** and females **(B)** at the acute time point. IL-5 levels increased in the periphery of LPS treated males at the chronic time point **(C)** but did not significantly alter circulating levels of females **(D)**. Circulating levels of IL-10 were increased at all time points in the males **(E)**, with LPS increasing IL-10 levels at the acute, chronic, and terminal time points. Moreover, at the acute time point, the combination of LPS and chronic stress decreased circulating IL-10 levels when compared to males that were treated with LPS alone. **F)** LPS increased circulating IL-10 levels in females at both the acute and chronic time points. Reported values depict mean ± SEM. *p < 0.05, **p < 0.01, ***p < 0.001, ****p < 0.0001.

IL-5 levels in males display a main effect of time point (F_(2,76)_ = 4.222, p = 0.0182), treatment (F_(1,76)_ = 7.609, p = 0.0073), and a time point by treatment interaction (F_(2,76)_ = 3.522, p = 0.0345; **FIGURE 9C**), driven by a significant difference between the saline and LPS treated males at the chronic time point (p = 0.0007). Female data only show a main effect of time point (F_(2,78)_ = 9.038, p = 0.0003; **FIGURE 9D**), with levels of IL-5 decreasing over the course of the injection series.

Data from circulating IL-10 levels show a complex set of interactions within the males. IL-10 levels showed a main effect of time point (F_(2,77)_ = 25.88, p < 0.0001), treatment (F_(1,77)_ = 63.84, p < 0.0001), stress (F_(1,77)_ = 5.124, p = 0.0264), as well as interactions between time point and treatment (F_(2,77)_ = 22.79, p < 0.0001), time point and stress (F_(2,77)_ = 6.060, p = 0.0036), treatment and stress (F_(1,77)_ = 4.081, p = 0.0469), and treatment by time point by stress (F_(2,77)_ = 5.357, p = 0.0066; **FIGURE 9E**). Post hoc analysis showed a significant difference between the non-stress groups (p < 0.0001), the LPS groups (p = 0.0014), and stress groups (p = 0.0154) at the acute time point. Moreover, there is a treatment effect at the chronic (p < 0.0001) and terminal (p = 0.0079) time points. Collectively, LPS treatment elevates IL-10 after acute and chronic dosing and it remains elevated for at least a week past the final exposure. Female data show a main effect of time point (F_(2,78)_ = 25.12, p < 0.0001) and treatment (F_(1,78)_ = 59.51, p < 0.0001; **FIGURE 9F**). Data also display a time point by treatment interaction (F_(2,78)_ = 24.39, p < 0.0001), which is driven by a significant difference between saline and LPS treated females at the acute (p < 0.0001) and chronic (p = 0.0008) time points.

##### 2.16 Circulating ROMO-1Cconcentration

Males exposed to chronic injections of LPS displayed increased circulating ROMO1 than their saline treated male counterparts regardless of stress history (F(_1,27_) = 4.984, p = 0.0341; **FIGURE 10A**). Despite a visual difference in means, a history of chronic stress did not significantly elevate ROMO-1 concentrations (p > 0.05). Within the female groups, there was no significant difference between stress or treatment background (p > 0.05; **FIGURE 10B**).

**Figure 10:**
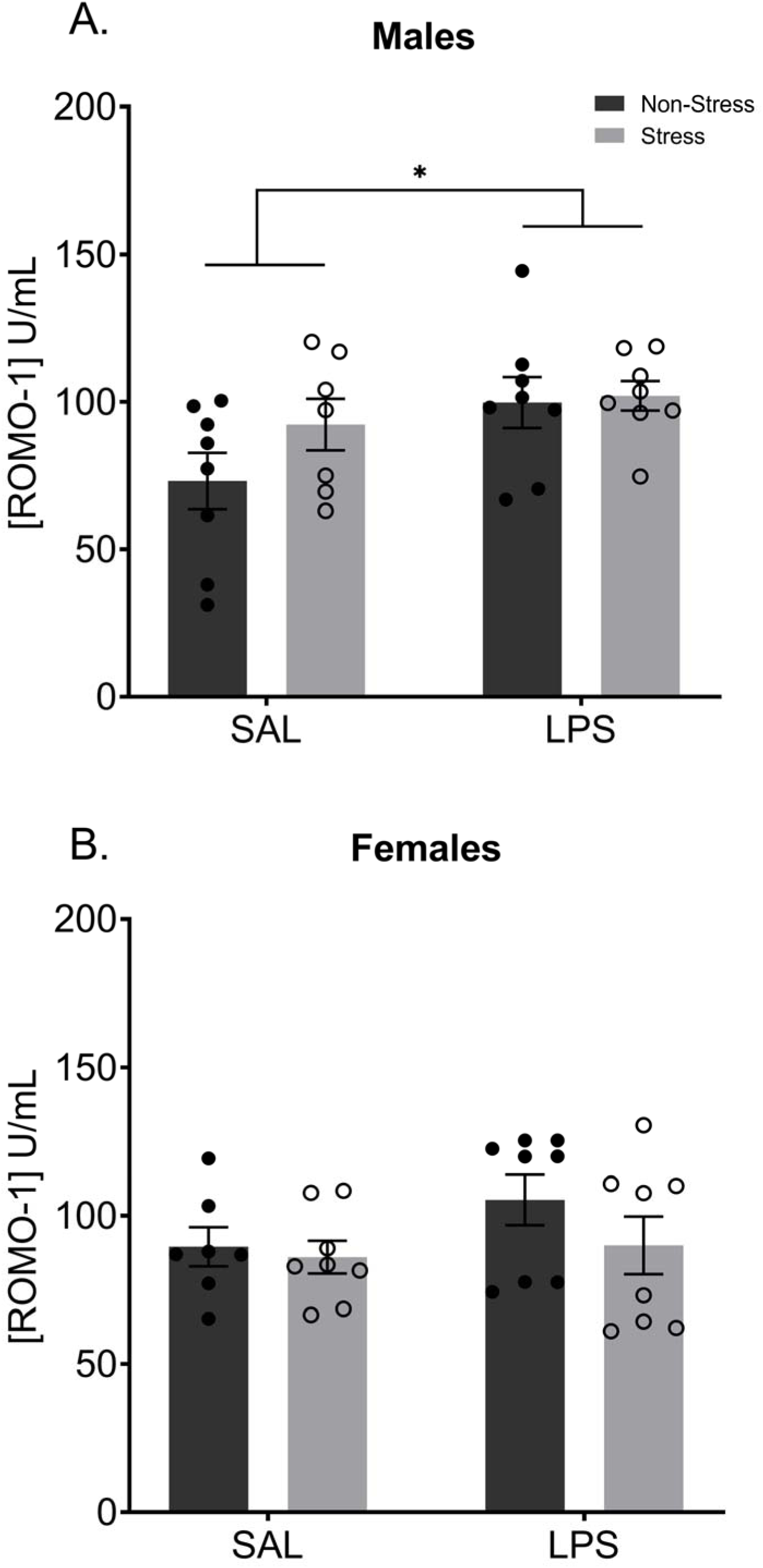
Chronic LPS Increases Circulating ROMO-1 in Males. Terminal levels of peripheral ROMO1, eleven weeks after the final LPS treatment, were significantly increased in males treated with LPS (**A**) but not females (**B**). There was no effect of stress history on circulating levels of ROMO1 at baseline in either males or females. Reported values depict mean ± SEM. *p < 0.05.

## Discussion

A history of chronic traumatic stress is sufficient to alter mitochondrial function in synaptosomes from female mice three weeks after exposure to stress. In addition, with the exception of indicators of increased anxiety-like behavior, there were no observed phenotypic indicators of the neural changes. In contrast, males were susceptible to only inflammation-induced reductions in synaptic respiration as well as effects of chronic stress and chronic inflammation on piloerection and inflammatory profiles both peripherally and centrally. These data suggest that female dominant manifestation of neurodegenerative conditions such as Alzheimer’s disease (Ferretti et al., 2018) may in part be fueled by invisible damage accrued by life stressors and inflammatory challenges; whereas, males are more likely to manifest observable shifts in inflammatory metrics.

Despite elevated synaptosomal respiration in females with a history of chronic stress or chronic inflammation, there was a reduced number of mitochondria present at the synapse in synaptosomes of females with a history of chronic LPS. This reduction suggests an impaired rate of mitochondrial trafficking to the synaptic space (Fang et al., 2012). Combined with increased respiration, it is possible that females who have experienced chronic inflammation or a history of stress have increased individual mitochondrial respiration, potentially compensating for the deficit of mitochondrial number at the synapse via increased mitochondrial productivity. Compensatory mechanisms for the individual exposure to either chronic life stress or chronic inflammation appear to wane when exposed to both conditions such that in females that experienced chronic inflammation after a history of chronic stress, synaptosomal respiration was decreased. In particular, females with both a history of stress and chronic inflammation displayed reduced proton leak, therefore, decreased respiration is likely due to a deficit in the oxidative phosphorylation pathway (Cheng et al., 2017; Krauss et al., 2002). Proton leak, the connecting factor between oxygen consumption and ATP generation, does not work alone and is minimally assisted by the ‘electron slip’ that may give rise to an elevated oxygen consumption rate disproportionate to what would be expected (Cheng et al., 2017), thus acting as a compensatory mechanism in times of mitochondrial distress. This shift in mitochondrial function likely prevents the activation of this protective compensatory mechanism, leaving ATP production at the synapse compromised when chronic inflammation is combined with chronic stress. This decreased respiration could lead to impaired neurotransmission, as ATP is critical in neurotransmitter packaging and vesicular release at the presynaptic terminal (Pathak et al., 2015; Rangaraju et al., 2014). Taken together, our data suggest that females employ compensatory mechanisms when exposed to a single modality of chronic exposure, stress or inflammation. However, when a consummate capacity is reached, as observed here in the females with a history of chronic stress combined with chronic inflammation, these mechanisms become inaccessible and synaptic respiration is compromised possibly leading to eventual compromise of neural function. Finally, females did not exhibit evidence of a proinflammatory neural environment, as indicated by the cytokines assessed, despite evident peripheral effects of initial LPS exposure. This may be indicative of an uncoupling of inflammation and metabolism in the female brain, as compared to the male brain, and should be a course of future study.

The inflammatory function of males appears more labile and influential at the neural level than in females, such that metabolic data show that chronic inflammation alone is enough to decrease synaptic respiration. Interestingly, this effect compounds in males with a history of stress suggesting that stress creates a more vulnerable respiratory neural environment to the effects of inflammation. We did not observe changes in physical characteristics of mitochondrial structure or synaptic enrichment in male samples which suggests that effects on respiration were the product of an overall decrease in mitochondrial function via a mechanism not directly related to mitochondrial health. Potential explanations for this phenomenon could lie in increased rate of mitophagy (Wang et al., 2019) or a decrease in the mitochondrial proton motive force in a mechanism not directly assessed through our analysis (Berry et al., 2018). Moreover, the null results in central ROMO1 expression data suggest that mitochondrial respiration differences are not due to altered levels of reactive oxygen species in the brain, but the elevated ROMO1 concentrations in the blood may implicate peripheral effects of reactive oxygen which were not detectable at the expression level in the brain.

Inflammation could be a source of altered synaptosome mitochondrial in males given that they display increased central and peripheral impacts of stress and chronic LPS exposure. Hippocampal expression of the pro-inflammatory cytokines TNF-α and IL-1ß were increased following chronic inflammation – an effect exacerbated by stress history on TNF-α levels. This is particularly remarkable because the last predatory stressor was eleven weeks prior to sample collection and the last LPS exposure was one week prior to collection. Males also showed an increase in pro- (IL-6, KC-GRO) and anti- (IL-5, IL-10) inflammatory peripheral cytokines after both the initial and final chronic injection of LPS. These effects are to be expected, as an initial immune challenge promotes the production and circulation of systemic inflammation in response to a challenge (Couper et al., 2008; Deslauriers et al., 2017; Pacholko et al., 2019; Pripp and Stanišić, 2014); however, the observed differences at the terminal endpoint suggest a more sustained impact on inflammatory regulation following chronic stress and chronic inflammation. Taken together, these data suggest that males are susceptible to the effects of chronic inflammation and that a history of stress exacerbates this vulnerability at the level of whole brain synaptic respiration and the pro-inflammatory response.

Our results provide evidence that chronic repeated predation stress beginning in adolescence is sufficient to produce an anxiety-like phenotype, independent of sex, that is observable long after removal from the stressful environment. Accompanying this anxiety-like phenotype, we observed sex-specific changes in mitochondrial bioenergetics that highlight a potential mechanism by which stress and inflammation, either alone or together, can modify synaptic function. These data further support the considerable evidence that males are more susceptible to the negative effects of chronic inflammation than females (Bekhbat et al., 2019) and that females require compounded insults before inflammation-mediated deficits manifest. Our results expand the growing knowledge relating to the interplay of stress, inflammation, and sex and promote mitochondrial modulation as a bridging factor between chronic stress, chronic inflammation, and neuropsychiatric disease. Future research is needed to uncover the specific mitochondrial components, the impact of altered respiration on synaptic function, and relevant compensatory pathways giving rise to the sex-specific effects observed in this study.

## Supporting information

Supplemental Methods & Results

## 5. Acknowledgements

The authors would like to thank the Division of Animal Resources at Virginia Commonwealth University (VCU) for their assistance with animal care. Microscopy was performed at the VCU Microscopy Facility, supported, in part, by funding from NIH-NCI Cancer Center Support Grant P30 CA016059. This work was supported by an Alzheimer’s Research Supplement to an R01 from National Institutes of Health National Institute of Nursing Research (NR014886; GNN). In addition, some funding for trainees was provided by NIH IRACDA Grant K12GM093857 (MMH) and an NIH IMSD Grant 5R25GM090084-08 (GAS).

