## Supplemental Methods & Results for "Traumatic stress history interacts with chronic peripheral inflammation to alter mitochondrial function of synaptosomes in a sex-specific manner"

### 1.0 Supplemental Methods

#### *1.1. Chronic Repeated Predatory Stress (CRPS)*

Mice chosen to undergo CRPS were isolate housed and deprived of nestlet enrichment beginning on PND 34 and throughout stress. Immediately following isolation and removal of enrichment, CRPS mice began exposure to daily predation stress, which involved protected exposure to an adult male Long Evans rat. Mice were placed in a clean dwarf hamster ball (Lee's Aquarium & Pet Products, San Marcos, CA, USA, Cat. #20198) and subsequently placed in the home cage of the predator, who were all adult male Long Evans rats previously used for breeding. Non-stress mice remained pair housed with enrichment and were handled daily. After the final stressor exposure on PND 71, CRPS mice received nesting material for the remainder of the study. Additionally, the non-stress mice were isolate housed on PND 71 for the remainder of the study to control for the effects of isolation housing on stress and sickness behavior (Eisenberger et al., 2016; Hennessy et al., 2013).

#### *1.2. Social Interaction*

The three-chambered sociability test was used to assess social anxiety-like behavior (Farrell et al., 2016; Moy et al., 2004; Nadler et al., 2004) following CRPS and chronic low-level inflammation. Social interaction testing occurred the morning before the 15$^{th}$ injection (PND 118), 72 hours after the 14$^{th}$ injection. Testing occurred in two phases: a five-minute habituation phase and a ten-minute testing phase. During the habituation phase, mice were allowed access to the middle chamber of a three-chambered apparatus (Med Associates, Fairfax, VT) with Plexiglas doors blocking the ability to enter the two adjacent chambers. After habituation, mice were returned to their home cage for an inter-trial interval of three minutes, where the chamber was cleaned with 70% ethanol and Plexiglas doors removed. A novel, sex- and age-matched conspecific was placed in a cup within one of the side chambers and an empty cup placed in the other (sides were counterbalanced across groups). Mice were allowed to explore all three compartments of the chamber for a total of ten minutes. Time spent in each of the three chambers, time and frequency interacting with the novel mouse, velocity, and distance traveled were assessed via overhead video and EthoVision 14.0 software. Behavioral data from social interaction were analyzed using two-way analysis of variance (ANOVA) with the factors of stress and LPS.

#### *1.3. Barnes Maze*

The Barnes Maze test was used to assess spatial learning and memory. The Barnes Maze consisted of a white, circular table (95cm high and 92 cm in diameter) with 20 equidistant open holes (five-cm in diameter) located on the perimeter of the maze in a room lit to 2500lux. Initial acquisition occurred twice a day for eight consecutive days beginning on PND 125. It is important to note that this was 4 days removed from the final LPS injection in the chronic series and physical assessments indicated that mice were not demonstrating overt sickness behavior by this timepoint (Figure 1). Mice were placed on the maze and given three minutes to locate the escape hole, which contained a nesting box in lieu of an open hole. Once the mouse entered the hole, the lights were turned off and a cover was placed over the hole to the nesting chamber. The mouse remained in the covered chamber for two minutes before being returned to its home cage for a three minute inter-trial interval. The maze and escape box were cleaned with 70% ethanol between each trial. The day following the final acquisition day each mouse was subject to a five-minute probe trial to assess memory where the escape box was removed and the latency to the goal location was assessed. Latency to reach the escape box, distance traveled, and velocity were assessed using an overhead camera and EthoVisionXT 14.0.

Reversal learning began 24 hours following the acquisition probe. Reversal training consisted of the goal box location being moved to the opposite quadrant from the location learned during Acquisition. Reversal training was conducted twice a day for five days. Memory for the new goal box location was conducted 24 hours after the final reversal learning day. A challenge injection was given 24 hours later and then a final probe was conducted three days later to investigate if an additional injection (SAL or LPS where appropriate) altered memory performance. Latency to reach the escape box, distance traveled, and velocity were assessed using an overhead camera and EthoVisionXT 14.0. Behavioral data from the Barnes maze probe trials were analyzed using two-way analysis of variance (ANOVA) with the factors of stress and LPS. Barnes Maze acquisition data was analyzed using a three-way repeated measures ANOVA with the factors of stress, LPS, and trial number.

#### *1.4. Tissue Collection*

Immediately following decapitation brains were extracted on ice and rinsed with isotonic sucrose solution (0.32 M sucrose, 1mM EDTA, 5mM Tris, pH 7.4) to remove excess blood. Brains were then bisected along the midsagittal plane. The hippocampus and prefrontal cortex were dissected from the right hemisphere and flash frozen on dry ice and stored at -80°C. The remainder of the right hemisphere and the entire left hemisphere were diced and homogenized in a seven-mL glass Dounce Homogenizer in isotonic homogenizing buffer (five up-and-down strokes of the loose plunger followed by five up-and-down strokes of the tight plunger) in preparation for synaptosomal isolation (concentrations for 'homogenizing buffer' and subsequent Percoll layers can be found in (Dunkley et al., 2008)).

### *1.5. Synaptosomal Isolation*

Synaptosomes isolation was adapted from (Dunkley et al., 2008). After the initial homogenization, samples were spun at 3600 RPM at 4°C for ten-minutes. The supernatant (approximately four-mL) was transferred to a clean, sterile 15mL conical tube and diluted to seven-mL using homogenizing buffer. Six-mL of the diluted supernatant was then carefully laid atop a five-layer discontinuous Percoll gradient in a 26mL polycarbonate ultracentrifuge tube. Gradients were spun at 32500xg for 15 minutes to separate synaptosomes from other neural fragments. Synaptosomes were extracted from the interface between the 15% and 23% Percoll layers and placed in a clean polycarbonate tube with 20mL of ionic media (20mM HEPES, 10mM D-Glucose, 1.3mM $Na_2HPo_4$, 1mM $MgCl_2$, 5mM $NaHCO_3$, 5mM KCl, 140 mM NaCl, pH 7.4; adopted from (Choi et al., 2009)) and spun for 35 min at 15000xg. Synaptosomal pellets were collected and measured via Nanodrop for protein content. Synaptosomal pellets were diluted to 40µg/100µL in Ionic Media in a new microcentrifuge tube. Remaining synaptosome pellets were saved at -20°C for Western blots or pelleted and stored in glutaraldehyde for imaging analysis. 100µL of diluted synaptosomes were plated in triplicate on a Seahorse XFe24 cell plate (Agilent Technologies, Santa Clara, CA) coated with poly-D lysine. The plate was spun for 30 minutes at 3400 RPM at 4°C to allow synaptosomes to adhere to the bottom of the well. Ionic media was aspirated out of each well and replaced with 500µL of freshly made assay media (prepared according to the manufacturer's instructions). Plates were incubated at 37°C in a non-$CO_2$ incubator for 30 minutes before measurement in the Seahorse XFe24 Analyzer (Agilent Technologies).

*1.6. Western Blotting*

Samples were pooled according to group to assure adequate protein amounts for loading. 20µg of protein, assessed via BCA assay (ThermoFisher Scientific, Cat. # 23225), was loaded and run using a Criterion XT 10% Bis-Tris midi gel (BioRad Laboratories, Hercules, CA, USA; Cat. # 3450113) and XT-MOPS running buffer (BioRad Laborotories, Cat. # 161-0788). Protein was transferred onto a PVDF midi membrane (BioRad Laborotories) using the TransBlot Turbo (BioRad Laboratories) system for dry transfers. Total protein levels within the gel were assessed using Revert 700 Total Protein stain (LI-COR Biosciences, Lincoln, NE, USA; Cat. # 926-11010) to normalize the gel and immediately imaged at 700nm using a LI-COR Odyssey system. The membrane was cut at 70kDa and 38kDa to allow separate incubation of each target. Total protein stain was stripped off of the membrane according to manufacturer's instructions and membrane was blocked with LI-COR Intercept Blocking Buffer (LI-COR Biosciences) for 40 minutes. Primary antibody was added and incubated at 4°C overnight on a shaker. Secondary antibody was added (LI-COR Biosciences; 800CW Goat anti-mouse $IgG_1$ Cat. # 926-32350; 680LT Donkey anti-goat IgG Cat. # 926-68024) and incubated at room temperature for 90 minutes on a shaker. Membranes were imaged on a LI-COR Odyssey Imaging system (LI-COR Biosciences) and analyzed using Image Studio Lite Version 5.2 (LI-COR Biosciences).

*1.7. Transmission Electron Microscopy*

To confirm sample enrichment and synaptosomal integrity, remaining pellets from the final ultracentrifugation step of the synaptosomal isolation were transferred to a 1.5mL microcentrifuge tube and spun at 4°C for ten-minutes at max speed. Supernatant was removed and pellets spun a second time to assure all remaining ionic media was removed. Pellets were then resuspended with 500mL 2% glutaraldehyde in 0.1M sodium cacodylate buffer with 0.1M sucrose at room temperature and allowed to sit undisturbed for ten-minutes. Samples were then spun at 4°C for ten-minutes at max speed to create a synaptosomal pellet. Pelleted samples were stored at 4°C until further processing.

Samples were rinsed in 0.1M cacodylate buffer a total of three times each. 2% osmium tetroxide in 0.1M cacodylate buffer was added as fixative and incubated for one-hour. Samples were rinsed again in 0.1M cacodylate buffer a total of three times. Fixed pellets were subsequently dehydrated in a graded ethanol series for five to ten minutes each, followed by three changes of 100% ethanol incubations for five to ten minutes each. Propylene oxide was then added in three successive changes followed by a 50/50 mix of propylene oxide and PolyBed 812 resin (Polysciences, Inc., Warrington, PA, USA, Cat. # 08791-500) in an overnight incubation. PolyBed 812 resin was then added in embedding molds and placed in a 60°C oven overnight. Embedded synaptosomal pellets were then sectioned at 700nm on a Leica EM UC6i ultramicrotome (Leica Microsystems, Buffalo Grove, IL, USA). Sections were subsequently stained with 5% uranyl acetate and Reynold's lead citrate.

Samples were imaged on a JEM-1400Plus transmission electron microscope (JEOL USA, Peabody, MA, USA) with a Gatan UltraScan 4000SP 4k x 4k CCD camera (Gatan, Inc., Pleasanton, CA, USA). Representative images of synaptosomes were obtained at a voltage of 100kV using a 15,000x magnification and 2.0 second exposure time. Images were obtained at a voltage of 100kV using a 4000x magnification and 2.0 second exposure.

### 2.0 Supplemental Results

#### *2.1 Social Interaction*

Using a two-way ANOVA with the factors of stress and treatment within each sex, data show no significant difference in the time spent in the chamber with the novel mouse ($p > 0.05$) in both males and females (**Supplemental Figure 1**). As an additional check, time spent interacting with the novel mouse, defined as the total time the nose point is in a predefined "cup zone", show no significant differences in either males or females ($p > 0.05$; data not shown). These findings suggest chronic repeated predation stress does not induce social anxiety-like behaviors, as shown in (Shaw et al., 2019a).

#### *2.2 Barnes Maze*

*Acquisition*

Analysis of acquisition data using a three-way repeated measures ANOVA with the factors trial, stress, and treatment show a main effect of trial within the males ($F_{(4.965,134.1)} = 116.0$, $p < 0.0001$; **Supplemental Figure 2A**) suggesting all males, regardless of group, learned where the goal box was by the end of the acquisition phase. There were no significant differences within the males when assessed in the memory probe ($p > 0.05$) when analyzed using a two-way ANOVA, suggesting no deficits in memory. Within the females, analysis of the acquisition data show a main effect of trial ($F_{(4.475,125.3)} = 99.36$, $p < 0.0001$; **Supplemental Figure 2B**) with a treatment by trial interaction ($F_{(7,196)} = 2.563$, $p < 0.0151$). Post hoc analysis suggests this interaction is driven by significant differences between saline treated and LPS treated female mice on acquisition days seven ($p = 0.0445$) and eight ($p = 0.0168$), with LPS treated females exhibiting increased total latencies to reach the goal box than their saline treated counterparts regardless of stress history. Similar to the males, there was no difference in the memory probe within the female mice ($p > 0.05$).

*Reversal*

Within the males, analysis of reversal data via three-way repeated measures ANOVA with the factors of time, stress, and treatment show a main effect of trial ($F_{(3.013, 81.36)} = 67.82$, $p < 0.0001$) and an interaction between stress and treatment ($F_{(1,27)} = 4.768$, $p = 0.0379$; **Supplemental Figure 2C**). Post hoc analysis show this interaction is driven by a significant difference in summed latency to the reversal goal box between non-stress saline treated males and non-stress LPS treated males ($p = 0.0425$). Female reversal data display a main effect of trial ($F_{(2.551, 71.43)} = 86.92$, $p < 0.0001$; **Supplemental Figure 2D**). There were no significant differences in the reversal probe ($p > 0.05$) within the female mice.

**Supplementary Table 1: Resulting Data for Levels of Circulating IL12p70 were Inconclusive.**

Majority of the samples analyzed fell below the curve, disabling our ability to perform statistical analysis on these samples. The number of samples that fell within the curve and their averages are noted in the table as “N” and “Mean” in pg/mL respectively.

| | | [redacted] | | |
|---|---|---|---|---|
| ***Males*** | Non-Stress SAL | N=1<br>Mean = 719.79 pg/mL | N = 1<br>Mean = 822.64 pg/mL | N = 1<br>Mean = 53.94 pg/mL |
| | Stress SAL | N = 2<br>Mean = 146.91 pg/mL | N = 1<br>Mean = 235.92 pg/mL | N = 2<br>Mean = 34.41 pg/mL |
| | Non-Stress LPS | N = 7<br>Mean = 1105.20 pg/mL | N = 2<br>Mean = 553.31 pg/mL | N = 3<br>Mean = 83.77 pg/mL |
| | Stress LPS | N = 5<br>Mean = 1278.81 pg/mL | N = 1<br>Mean = 630.96 pg/mL | N = 3<br>Mean = 64.43 pg/mL |
| ***Females*** | Non-Stress SAL | N = 4<br>Mean = 131.32 pg/mL | N = 4<br>Mean = 208.78 pg/mL | N = 4<br>Mean = 41.90 pg/mL |
| | Stress SAL | N = 3<br>Mean = 247.47 pg/mL | N = 3<br>Mean = 220.84 pg/mL | N = 6<br>Mean = 44.39 pg/mL |
| | Non-Stress LPS | N = 9<br>Mean = 1426.58 pg/mL | N = 0 | N = 1<br>Mean = 76.83 pg/mL |
| | Stress LPS | N = 7<br>Mean = 1041.10 pg/mL | N = 1<br>Mean = 313.08 pg/mL | N = 4<br>Mean = 24.33 pg/mL |

**Supplemental Figure 1.**

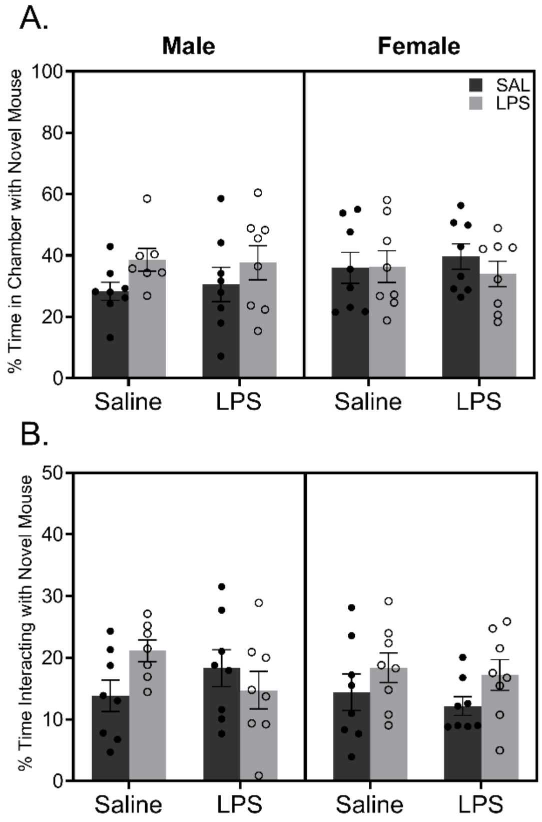


**Supplementary Figure 1: Neither a Chronic Predation Stress nor Chronic Inflammation Induced Changes in Sociability.**

An assessment of sociability was measured the morning before the fifteenth saline (SAL) or LPS injection. (**A**) Data reveal no significant difference in the percent time spent in the chamber containing the novel mouse ($p > 0.05$). **(B)** Additionally, there were no significant differences in the percent of time spent directly interacting with the novel mouse regardless of stress or treatment history.

**Supplemental Figure 2.**

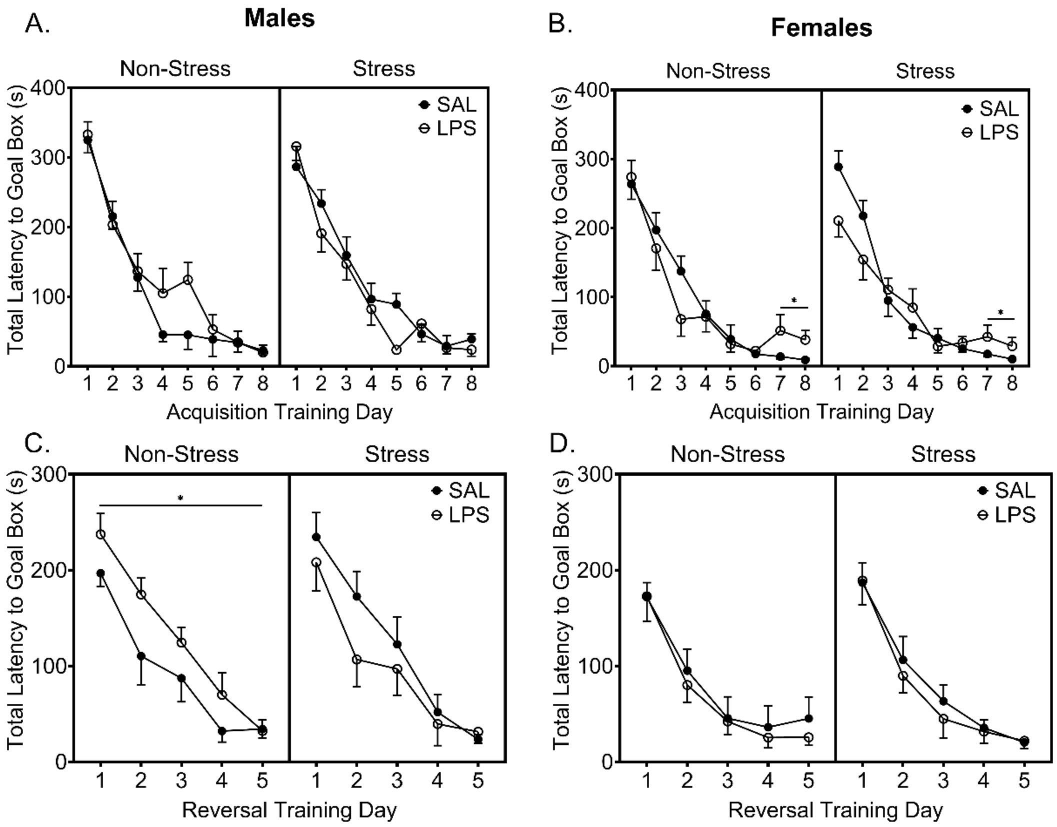


**Supplemental Figure 2: Chronic Low-level Inflammation Decreases Cognitive Flexibility in Male Mice with No Stress History. A)** In males, data show no difference by stress or treatment in latency to reach the goal box across the two trials per day suggesting no learning deficits between the groups **B)** Within the female mice, there was a significant increase in latency to reach the goal box in the LPS treated females regardless of stress history on days seven and eight. **C)** Male reversal data show a stress by treatment interaction ($F_{(1,24)} = 4.768$, $p = 0.0379$), with post hoc analysis revealing a significant increase in latency in the non-stress LPS group when compared to the non-stress saline treated controls ($p = 0.0425$), suggesting a decrease in cognitive flexibility **D)** There were no significant treatment or stress effects in the female reversal data. Reported values depict mean ± SEM. $*p < 0.05$.

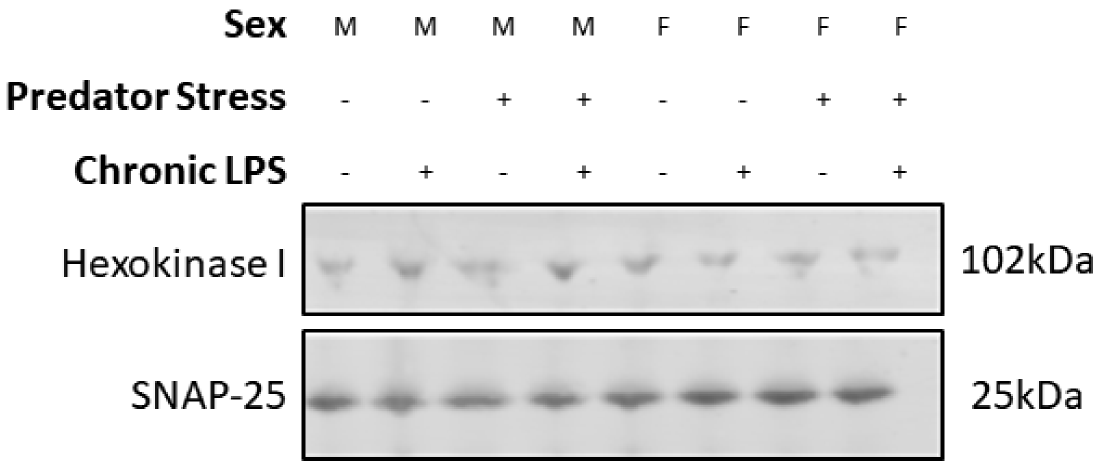


**Supplementary Figure 3: Western Blot Analysis of Synaptosomal Isolates.**

Synaptosomal isolates were pooled by group and probed for Hexokinase I and SNAP-25. Three technical replicates were completed for each group and used for analysis.
